# Oral LNAD+ Rapidly Elevates Intracellular NAD and Metabolic Flux Without Elevating Circulating NAD: Evidence from a Randomized Controlled Trial

**DOI:** 10.64898/2026.03.25.714130

**Authors:** Sergey A. Kornilov, Waylon J. Hastings, Lynne Fahey McGrath, Michael Leitz-Langan, Andrew T. Magis, Steve M. Coppess, Wendy Komac

## Abstract

Declines in nicotinamide adenine dinucleotide (NAD+) are linked to metabolic stress accompanying aging and disease. While precursor-based approaches elevate systemic NAD, their clinical translation can be constrained by biosynthetic bottlenecks and first-pass metabolism. RENEWAL-NAD+ (ClinicalTrials.gov NCT07336836; retrospectively registered 01/04/2026) was a double-blind, randomized, placebo-controlled Phase 0/1b trial in healthy adults aged 45–75 years (60 randomized; primary analysis n=50) evaluating 5 days of oral LathMized® NAD+ (LNAD+), a physicochemically modulated formulation that alters the supramolecular organization and solution behavior of NAD+ while preserving its native molecular structure. The primary endpoints were change in intracellular NAD (icNAD), measured in whole blood, and circulating NAD (cirNAD), measured in separated plasma, relative to baseline. LNAD+ produced a rapid and pronounced increase in icNAD, with a 53% elevation versus placebo at Day 6 (p=5.48e^−14^; Hedges’ g=3.66), while cirNAD was unchanged (p=0.60), demonstrating compartment-selective augmentation. Plasma NAD catabolites increased substantially (1-methyl-nicotinamide, MeNAM p=5.39e^−13^; N1-methyl-2-pyridone-5-carboxamide, 2PY p=2.95e^−16^), consistent with downstream engagement of NAD metabolic flux. Exploratory analyses identified non-overlapping correlates for the two compartments (cirNAD tracking inflammatory and metabolic markers, icNAD tracking red blood cell indices and NAM). Treatment was very well tolerated: symptom incidence was comparable between groups (p=0.68), only one mild adverse event (nausea, Grade 1) occurred in the LNAD+ arm, and no secondary clinical, vital-sign, wellbeing, or wearable-derived endpoint survived multiplicity correction. These data demonstrate rapid intracellular NAD augmentation after oral LNAD+ dosing with pharmacodynamic evidence of downstream metabolism, compartment-specific physiological signatures, and a favorable short-term safety profile, with exploratory multi-omic analyses ongoing.

## INTRODUCTION

Nicotinamide adenine dinucleotide (NAD+) is one of the most fundamental and evolutionarily conserved molecules in cellular metabolism. It serves dual roles as a coenzyme in over 500 enzymatic reactions and is a consumed substrate for several families of regulatory enzymes (1, 2). This pyridine nucleotide, discovered over a century ago by Harden and Young, is now recognized as a master metabolic regulator whose availability directly influences cellular energetics and organismal healthspan (3, 4). Thus, the importance of NAD+ in human physiology derives from its central role in bioenergetic pathways and extends to its recently described roles in epigenetic and post-translational regulation, circadian rhythm control, and intercellular communication (5, 6).

The centrality of NAD+ to the biology of aging has been reinforced by its mechanistic intersection with multiple hallmarks of aging (7, 8). As noted above, NAD+ serves as a co-substrate for enzymes including sirtuins (SIRT1–7), poly(ADP-ribose) polymerases (PARPs), and the cyclic ADP-ribose hydrolase CD38. Each of these consumes NAD+ stoichiometrically during catalysis. Age-related upregulation of CD38 expression is linked to chronic low-grade inflammation (“inflammaging”) and senescence-associated secretory phenotype (SASP) signaling, and is thought to be a strong driver of age-dependent tissue NAD+ depletion (9), creating a feed-forward cycle in which declining NAD+ impairs enzymes responsible for maintaining genomic stability (PARP1), mitochondrial quality control (SIRT1/SIRT3), and metabolic homeostasis (10, 11). This mechanistic convergence positions NAD+ repletion as a high-leverage point for interventional geroscience, consistent with a broad inter-tissue NAD regulatory principles described in the NAD World 3.0 model by Imai et al. (12).

Aging-accompanying immunosenescence is characterized by reduced adaptive immune reserve, impaired vaccine responsiveness, and altered leukocyte homeostasis. These phenotypes are tightly coupled to cellular bioenergetics and NAD+-dependent signaling. Because CD38 expression is enriched in immune compartments and increases with age, age-associated NAD+ depletion may disproportionately constrain immune metabolic fitness in older adults (13, 14). Recent evidence indeed suggests that NAD+ depletion in aged CD8+ T cells drives mitochondrial dysfunction and loss of stem-like properties. Importantly, these effects can be rescued via NAD+ repletion without inflammatory activation (13).

Given NAD+’s central role in cellular metabolism and regulation, alterations in NAD+ homeostasis are closely linked to human health and disease. Growing evidence from human studies and animal models suggests that tissue NAD+ levels decline with age, with reductions of 50–65% documented in multiple tissues by the eighth decade of life (15, 16). The clinical manifestations of NAD+ decline have been documented across diverse tissues and conditions, where plasma metabolomic analyses show dysregulation in both healthy aging and disease (17). Importantly, brain NAD+ levels measured *in vivo* also demonstrate significant age-related decline, with vulnerability in regions affected by neurodegenerative diseases such as Parkinson’s Disease and Alzheimer’s Disease (18–20). Skeletal muscle (a tissue with high metabolic demands and limited regenerative capacity in aging) exhibits pronounced NAD+ depletion in age-related sarcopenia and is hypothesized to contribute to both impaired mitochondrial function and reduced exercise capacity in aging (21). The clinical relevance of these findings encompasses a broad spectrum of age-related diseases where NAD+ depletion represents both a biomarker and pathogenic factor.

The recognition that NAD+ depletion represents a modifiable risk factor for disease has generated considerable interest in therapeutic strategies to restore NAD+ homeostasis. Current approaches predominantly rely on administration of NAD+ precursors including nicotinamide (NAM), nicotinic acid (NA), nicotinamide riboside (NR), and nicotinamide mononucleotide (NMN). While precursor-based strategies have demonstrated efficacy in elevating systemic NAD+ levels in human trials, fundamental limitations constrain their therapeutic efficacy and clinical translation.

Bioavailability cascades create sequential barriers that reduce effective delivery of oral NAD+ precursors to target tissues. Recent studies found that the gut microbiome plays a dominant and previously unappreciated role in precursor metabolism, with bacterial deamidases converting over 95% of orally administered NMN and 80–85% of NR to deamidated forms before systemic absorption (22, 23). Following initial absorption, hepatic first-pass metabolism presents the next barrier, with the liver expressing high levels of nicotinamide N-methyltransferase that rapidly clears NAM from circulation through methylation, creating a metabolic burden and potential depletion of S-adenosylmethionine pools (24).

Even when precursors successfully navigate metabolic barriers, tissue-specific heterogeneity in NAD+ responses impose additional constraints. Most human studies measure NAD+ in a single biological compartment, with the majority measuring whole-blood concentrations. While blood NAD+ levels reliably increase with precursor administration (ranging from approximately 22–370% with NR and 11–60% with NMN in clinical trials), translation to metabolically relevant tissues shows marked variability (25, 26). The disconnect between systemic and tissue NAD+ elevation is exemplified by studies that demonstrate high (e.g., +90%) blood NAD+ increases without corresponding muscle elevation in metabolically compromised individuals, suggesting factors such as transporter expression, local enzyme capacity, and tissue-specific NAD+ consumption regulate the therapeutic efficacy of precursor-based NAD+ repletion (27–30).

Considering these limitations, direct NAD+ administration has long been recognized as theoretically ideal but practically challenging due to the molecule’s physicochemical properties. Evidence reviewed in a recent report prepared by the University of Maryland regarding inclusion of NAD+ in the 503B Bulks List suggested that current NAD formulations were susceptible to degradation and not stable as oral formulations. Nonetheless, it is overall supportive of positive therapeutic effects in human studies that administered exogenous NAD+ or NADH in patients with fatigue, Parkinson’s Disease, and addiction (31). NAD+ has a molecular weight of 663.43 Daltons, exceeding typical absorption thresholds, multiple negatively charged phosphate groups confer extreme hydrophilicity at physiological pH (logP = −11.6); its lability directly translates into poor shelf stability; and its susceptibility to rapid enzymatic degradation in the gastrointestinal tract limits half-life to less than five minutes (32, 33). These characteristics have historically led to oral NAD+ bioavailability estimates placing below 2%, thereby removing this route as a viable therapeutic option for NAD+ repletion.

Intravenous (IV) NAD+ administration is aimed at enhancing bioavailability but presents its own significant limitations. Hawkins et al. demonstrated that 500 mg intravenous NAD+ produced only a 2% increase in whole blood NAD+ at 24 hours, levels which were substantially inferior to both IV and even orally administered NR (34). Moreover, the rapid systemic clearance (half-life ∼1.7 hours), requirement for prolonged infusions, substantial incidence of adverse events including flushing and chest discomfort, and impracticality for chronic administration limit the utility of IV NAD+.

The development of LathMized® NAD+ (LNAD+) therefore represents a systematic response to the biochemical barriers limiting the effective oral administration of NAD+. The proprietary LathMize® platform was developed by Bryleos (Johnson City, TN) as a green chemistry and physical pharmacy process enabling solventless manufacturing of fragile endogenous molecules without altering their chemical identity. LNAD+ was engineered to enhance physicochemical stability and support systemic and intracellular availability following oral administration.

The RENEWAL-NAD+ study presented in this report was motivated by the dearth of evidence on oral NAD+ bioavailability and limited characterization of the mechanisms of action of NAD+ repletion. This study incorporated compartmentalized measurements of NAD+ in plasma (circulating NAD, cirNAD) and in whole blood (intracellular NAD, icNAD), allowing assessment of the relationship between systemic NAD+ pools and intracellular NAD+ dynamics. The majority of human NAD+ supplementation studies report whole-blood NAD+ concentrations, which predominantly reflect erythrocyte NAD+ pools without distinguishing between circulating and cell-associated fractions. This lack of compartmental resolution has been recognized as a barrier to interpreting the translational significance of NAD+-boosting interventions (35, 36), particularly relevant in aging research, where interventions may differentially affect extracellular pools, erythrocytes, and immune cell compartments, complicating cross-trial comparisons that rely solely on whole-blood measurements. In this study, icNAD reflects a whole-blood, predominantly erythrocyte derived intracellular component rather than a tissue or a measure specific to peripheral mononuclear cells (PBMCs).

The present first-in-humans study was therefore designed as a key translational step in establishing the safety, tolerability, and bioavailability of LNAD+ in healthy aging adults. The study had three primary objectives: (1) to determine whether oral LNAD+ administration increases intracellular NAD+ concentrations as measured in the red blood cells in whole blood vs. circulating NAD+ as measured in separated plasma alone; (2) to characterize the metabolic fate of administered NAD+ through quantification of downstream catabolites in plasma; and (3) to establish the safety profile of short-term oral NAD+ supplementation across clinical laboratory panels, vital signs, patient-reported outcomes, and wearable-derived activity and sleep measures.

## METHODS

### Study Design, Randomization, and Intervention

RENEWAL-NAD+ was a double-blind, randomized, placebo-controlled study (ClinicalTrials.gov NCT07336836; retrospectively registered 01/04/2026) to evaluate the safety, tolerability, and pharmacodynamic effects of five days of oral LNAD+ in healthy adults aged 45 to 75 years. ISB BioAnalytica Inc. (Seattle, WA) functioned as the Contract Research Organization, responsible for study design, implementation, and operational oversight in accordance with ICH GCP guidelines. The Institute for Systems Biology (ISB) provided scientific collaboration and bioanalytical expertise. The trial was retrospectively registered on ClinicalTrials.gov (NCT07336836) due to administrative delay; primary and secondary endpoints were specified in the statistical analysis plan prior to database lock and unblinding.

Inclusion criteria required participants to abstain from confounding supplements for a seven-day washout period. Participants had to be non-pregnant with no history of chronic or acute infections within the last two months and have BMI < 35 kg/m^2^. Additional exclusion criteria included current cancer or hematologic conditions, Type I or Type II diabetes, inflammatory bowel disease, autoimmune disorders, and use of systemic anti-inflammatory or immunosuppressive medications, with the exception of NSAIDs and acetaminophen.

Treatment allocation into LNAD+ and placebo arms was performed at enrollment using stratified *a priori* blocked randomization schemes. Randomization schemes were derived for males and females separately using varying block lengths (between K = 2 and K = 4) using R package *blockrand* (v.1.5). If participants failed the pre-screening or contact was lost prior to the initiation of the study protocol, new participants were randomized into the freed slot. Biological sex was therefore always included in statistical analysis as a covariate due to stratification. Overall, the study recruited 76 individuals of which 60 were enrolled, randomized to treatment, received study kits, and initiated the protocol; 51 participants completed the washout period, received at least one treatment dose, and provided at least one on- or post-treatment blood draw (N = 23 in LNAD+; N = 28 in the placebo group; see **Supplementary Figure 1 & Supplementary Table 1**).

LNAD+ is an oral powder that was supplied by BioNADRx d.b.a Bryleos in opaque plastic bottles with a masked label. NAD+ constituted 50% of the total mass of powder, with the remainder being an inactive filler of polyethylene glycol (PEG), a widely used pharmaceutical excipient with a well-established safety profile; it is generally considered biologically inert, minimally absorbed when administered orally, and primarily associated with mild, transient gastrointestinal adverse effects at higher doses (37). Additional orthogonal physicochemical characterization of the LNAD formulation was conducted using complementary analytical approaches to assess elemental composition, molecular features, and solid-state structure. These analyses confirmed compositional consistency, molecular identity, and distinct structural attributes of the formulation. Detailed methods and results are not disclosed due to proprietary considerations. Placebo powder contained only PEG as an inactive ingredient (100% by mass).

Following a 7-day washout period, participants were administered 2000 mg LNAD+ or placebo for five days following a QID schedule, with four 500 mg daily doses spread approximately 3–4 hours apart within a 10–16-hour window. On Days 2 and 3 an additional dose of 500 mg was taken at nighttime. Treatments were delivered by oral administration as a swish-and-swallow protocol.

### Blood Collection

Venous blood specimens (2–4 mL) were collected into K -EDTA Vacutainer® tubes (BD Vacutainer®, Franklin Lakes, NJ) by certified phlebotomists during home visits. Samples intended for NAD+ measurement were immediately placed on ice packs within insulated containers and shipped overnight to Jinfiniti’s facility under cold-chain conditions (2–8°C), with transit times limited to 24–36 hours. Upon receipt, specimens were logged, inspected for hemolysis, and stored pending immediate processing. Additional blood samples received at ISB were processed in a BSL-2 facility to isolate plasma, aliquoted, frozen, and biobanked for storage and further processing. Two baseline blood samples were collected during the one-week washout period 3–4 days and 1–2 days prior to study onset. On-protocol blood draws were collected on Day 4 (after 3 days of treatment) and on Day 6 (one day after the final LNAD+ dose). The blood draw on Day 4 corresponds to a cumulative total of ∼14 doses (500mg each); the final blood draw on Day 6 is taken after the last dose for cumulative intake of ∼22 doses at 500mg each or 11g total NAD+ administered.

### Primary Endpoints: Intracellular and Circulating NAD+

Peripheral venous blood samples were analyzed for circulating and intracellular NAD by Jinfiniti Precision Medicine (Augusta, GA, USA), a laboratory specializing in NAD+ quantification that subsequently attained CLIA certification. Intracellular NAD (icNAD) was quantified in the whole blood. Half of the blood was centrifuged to separate plasma from the cellular fraction and used to measure circulating NAD (cirNAD). The distinction between these compartments is methodologically important: cirNAD reflects extracellular NAD+ pools, while icNAD predominantly reflects erythrocyte NAD+ content, providing a measure of cellular NAD+ status.

The Jinfiniti assay employs a proprietary enzymatic cycling assay to generate a colorimetric signal proportional to total NAD (NAD+ + NADH) concentration. The assay demonstrated an intra-assay coefficient of variation (CV) of 3.1% based on duplicate measurements within single runs (provided by Jinfiniti), with a linearity of 0.997 in the range of 0–120 μM and a lower limit of quantification of 46 nM. In the RENEWAL-NAD+ study, we directly estimated test-retest reliability across two baseline timepoints at r = 0.97 (p < 0.001) for icNAD and r = 0.79 (p < 0.001) for cirNAD, indicating excellent assay reproducibility.

### Safety Endpoints

Safety was evaluated using a combination of vitals, clinical laboratory values, and patient-reported outcomes. Vital signs included blood pressure and heart rate measured twice at each blood draw visit and supplemented by wearables data from Fitbit Luxe devices provided to participants, which also measured activity (steps) and sleep quality (time to fall asleep, sleep duration). Clinical laboratory values included complete blood count and blood chemistry 24 panels measured in a CLIA-certified laboratory (LabCorp). Patient-reported adverse event incidence was tabulated at the end of the study using data volunteered by the participants. Additional patient-reported outcomes included baseline and daily physical symptoms reported using the Review of Systems questionnaire (RoS), which captures binary data across 71 symptoms organized into 14 bodily systems, as well as measures of subjective well-being assessed using the Clinically Useful Anxiety Outcome Scale-Daily version (CUXOS-D) (38), the Depression, Anxiety, and Stress Scale-21 (DASS-21) (39), and Daily Fatigue Impact Scale (D-FIS) (40).

### Secondary Endpoints: NAD+ Metabolites

Plasma abundance of NAD+ metabolites was assayed by the Metabolon Global Platform and included measurement of 2PY, nicotinamide (NAM), methylated NAM (MeNAM), trigonelline, and quinolinate. Frozen aliquots with 200 μL of plasma were shipped on dry ice to Metabolon upon completion of the study and stored at −80°C prior to metabolomic assaying using previously described procedures (41). Briefly, after precipitation of proteins in methanol and centrifugation, supernatant evaporated from each sample was aliquoted into three portions to undergo ultra-high-performance liquid chromatography/tandem mass spectrometry (UHPLC/MS/MS) for detection of polar as well as nonpolar metabolites. Metabolite peak detection, quality control, and metabolite identification were performed using Metabolon’s proprietary software, controlling for day-to-day run and between-batch variation using internal and pooled technical replicate samples, as well as purified standards library. Metabolite abundance was estimated in Metabolon’s proprietary relative abundance units, which represent normalized peak intensities after Metabolon platform-specific scaling and batch normalization (41).

Additional plasma and blood biospecimens were collected for exploratory multi-omic analyses, including proteomic and metabolomic profiling; as these analyses were not pre-specified endpoints for RENWEWAL-NAD+, they are not reported here.

### Secondary Endpoints: Oxidative Stress

Measures of oxidative stress included serum gamma-glutamyl transferase (GGT) and reactive oxygen metabolites. Serum GGT was measured as part of the standard blood chemistry profile (LabCorp), and reactive oxygen metabolites (O_2_, O_2_•^-^, H_2_O_2_, OH•, & OH^-^) were assayed by Jinfiniti.

### Secondary Endpoints: Organ Function, Inflammation, and Metabolism

All secondary endpoints of organ function, inflammation, and metabolism were obtained from complete blood count and blood chemistry panels (LabCorp). Indicators of liver function included levels of alanine transaminase (ALT), aspartate transaminase (AST), alkaline phosphatase (ALP), and total bilirubin. Indicators of kidney function included serum levels of blood urea nitrogen (BUN), creatinine, and estimated glomerular filtration rate (eGFR). Metabolism was estimated via fasting glucose, serum triglycerides, low-density lipoprotein (LDL), and high-density lipoprotein (HDL). Inflammation was assessed via white blood cell counts, serum levels of high-sensitivity C-reactive protein (hs-CRP), tumor necrosis factor alpha (TNF-α), and interleukin-6 (IL-6).

### Power Calculations

The study sample size was originally determined by a preliminary analysis of statistical power carried out in GPower 3.0 (42) with the following assumptions: a one-tailed statistical test; Type I error rate (nominal, without correction for multiple biomarkers) was set at 5% one-sided, to arrive at an optimistic power estimate given the directional hypotheses testing planned initially. Power analysis was performed for the F family of tests for repeated measures ANOVAs (3 measurements). Expectations regarding observed effect sizes were guided by the general considerations of effect size distributions in biomarker studies as well as conventional categories for Cohen’s (1988) f^2^ classifying values above 0.10 as small, 0.25 as moderate, and 0.40 as large (43). Cohen’s f^2^ thresholds were used solely for sample size estimation in power analyses and are not used for interpretation of study results.

The primary parameter of interest was statistical significance of an interaction term between treatment Group × Time or a within-between interaction. At 30 total participants (15 per arm), the study was projected to attain power of ∼80% to detect very large effect sizes (local Cohen’s f > 0.50).

Increasing the total target sample to 60 (30 per arm) was determined to enable the study to consider effect sizes in the medium range, with power fluctuating from 50% for f = 0.25 to ∼80% for f = 0.35. The study anticipated an attrition rate of 20% and was not designed to accommodate corrections for multiple testing across secondary endpoints. The study targeted N=60.

### Statistical Analyses

All analyses were performed in R (version 4.4.2) using a reproducible {targets} pipeline with computational environment locked via renv. Primary and secondary endpoints were analyzed using Mixed Models for Repeated Measures (MMRM), implemented via the *mmrm* R package (version 0.3.11), which provides robust inference for longitudinal clinical trial data with missing values under the missing-at-random (MAR) assumption. The MMRM approach models within-subject correlation through the residual covariance structure rather than random effects, making it particularly well-suited for clinical trials with structured visit schedules.

The model was specified as: CHG = BASE + AVISIT × TRT01A + SEX + AGE + BMIBL + VITDFL + SEX:AGE + us(AVISIT | USUBJID), where CHG represents the change from baseline, BASE the baseline value (average of two baseline measurements for clinical data), AVISIT the categorical visit variable, TRT01A the treatment assignment, and covariates for sex, age, baseline BMI, and vitamin D supplementation status. An unstructured (US) covariance matrix was specified for the within-subject residuals, allowing each variance and covariance to be estimated freely. The Kenward–Roger approximation was employed for denominator degrees of freedom, providing appropriate small-sample inference. Continuous covariates (age, BMI) were centered at their mean values. Visit-specific treatment contrasts and p-values were obtained via the *emmeans* R package (v1.10), with two-sided 95% confidence intervals.

For the primary NAD endpoints, nonparametric bootstrap confidence intervals were implemented using bias-corrected and accelerated (BCa) methodology with subject-level case resampling. The pre-specified MMRM was fit to the complete data to obtain the point estimate, then refitted to bootstrap replicate datasets generated by resampling subjects with replacement. A parallel jackknife pass (successively leaving out one subject at a time) provided the acceleration correction. To address multiplicity across clinical laboratory panels, p-values underwent Benjamini–Hochberg false discovery rate correction within outcome families (e.g., liver function, lipid metabolism, hematology). For global multiplicity assessment across families, the effective number of independent tests (M_eff_) was estimated using the Galwey (44) eigenvalue method applied to a regularized Ledoit–Wolf shrinkage covariance matrix of test statistics, with Šidák correction applied at the M_eff_-adjusted significance threshold. Symptom incidence from the Review of Systems (RoS) questionnaire was analyzed using binomial generalized linear mixed models (GLMMs) as implemented in glmmTMB, with treatment × visit interaction as the fixed effect of interest and random intercepts for subjects. Additional details regarding data processing and quality control are described in **Supplementary Methods** and **Supplementary Tables 2-3**.

Analyses were conducted on two overlapping analysis sets: the full intent-to-treat population (N = 51) and a primary analysis excluding one participant in the placebo group (N = 50) whose icNAD levels demonstrated a substantial decline after study initiation, consistent with recent discontinuation of B-vitamin supplements (a protocol-endorsed exclusion criterion). This exclusion was made after consultation with a reviewer after the unblinding of treatment assignment. Results were materially unchanged between the two analysis sets, and the primary analysis cohort (N = 50) is presented throughout. Alignment between analyses within the full intent-to-treat population (N = 51) and analysis cohort (N = 50) are presented in **Supplementary Figure 2**. Effect sizes for results are interpreted in the context of conventional categories for Hedge’s g classifying values above 0.20 as small, 0.50 as moderate, and 0.80 as large (45).

### Changes to Trial Protocol-Statistical Analysis Plan

Five methodological refinements were implemented following the initial protocol and the statistical analysis plan (SAP). First, an additional baseline blood draw was incorporated to enable derivation of more reliable baseline measurements, thereby increasing statistical power for within-subject change-from-baseline analyses.

Second, the primary analytical framework was updated from mixed ANOVA/ANCOVA to MMRM. This transition reflects current best practice in longitudinal clinical trial analysis, as recommended by regulatory guidance (ICH E9(R1)), and offers superior handling of missing data under the MAR assumption, explicit modeling of within-subject correlation structures through the unstructured covariance specification, and direct baseline covariate adjustment. The MMRM approach is more conservative than the originally planned ANOVA in that it makes fewer distributional assumptions about the correlation structure while providing narrower confidence intervals through efficient use of all available data.

Third, the SAP evolved from one-tailed hypothesis testing to two-sided tests with corresponding two-sided 95% confidence intervals. This more conservative inferential standard was adopted to align with conventional reporting norms; all reported treatment effects (primary endpoints) remain highly significant under this more stringent criterion. Fourth, for the primary endpoints, nonparametric bias-corrected and accelerated (BCa) bootstrap confidence intervals were implemented using subject-level case resampling to complement parametric MMRM inference, providing robust evidence of treatment effects without reliance on distributional assumptions. Fifth, blood draw windows were expanded to allow collection on variable days, including expanding baseline blood draws 3-4 days and 1-2 days prior to study onset, as well as adapting on-protocol blood draws for one participant to be on Day 3 and Day 5 instead of Day 4 (on protocol draw #1) and Day 6 (one day following final LNAD+ dose).

Collectively, these refinements were judged to increase the conservatism of the analysis and inference relative to the original protocol and SAP. The primary results are robust to the choice of analytical framework: MMRM, the originally planned ANOVA, and nonparametric bootstrap all yield concordant conclusions regarding the primary endpoint.

## RESULTS

### Sample Demographics

Study demographics are presented in **Table 1**. The LNAD+ and the placebo arms did not differ at baseline with respect to any of the examined characteristics including clinical laboratory test values, frequency of reporting of medical conditions, or medications usage (all p’s > 0.05). In addition to self-reported data, plasma metabolomics were utilized to determine tobacco usage or nicotine exposure (i.e., smoking status). Tobacco metabolites were only detected in two participants in the placebo group; both were retained in analyses.

**Table 1.** Baseline demographics and characteristics.

| Category | Characteristic | LNAD+ (N=23) | Placebo (N=27) |
| --- | --- | --- | --- |
|  | N | 23 | 27 |
| Age (years) | range | 48.0 – 72.0 | 46.0 – 72.0 |
|  | mean (SD) | 57.3 (7.3) | 59.3 (7.2) |
| Sex, n (%) | Female | 12 (52.2%) | 14 (51.9%) |
|  | Male | 11 (47.8%) | 13 (48.1%) |
|  | Postmenopausal | 1 | 2 |
| **Race, n (%)** | White | 17 (73.9%) | 25 (92.6%) |
|  | Asian | 2 (8.7%) | 1 (3.7%) |
|  | Black or African American | 0 (0.0%) | 1 (3.7%) |
|  | Multiracial | 4 (17.4%) | 0 (0.0%) |
|  | Other race | 0 (0.0%) | 0 (0.0%) |
| **Ethnicity, n (%)** | Not Hispanic or Latino | 19 (82.6%) | 24 (88.9%) |
|  | Hispanic or Latino | 2 (8.7%) | 1 (3.7%) |
|  | Other/Unknown ethnicity | 2 (8.7%) | 2 (7.4%) |
| **Medical history, n** | Depression | 6 | 4 |
|  | Anxiety | 3 | 5 |
|  | Hypertension | 4 | 4 |
|  | Insomnia | 3 | 2 |
|  | Sleep apnea | 2 | 2 |
|  | Hyperlipidemia | 1 | 2 |
|  | Hypothyroidism | 2 | 1 |
|  | Heart attack | 1 | 0 |
| **Baseline vitals** | range | 19.6 – 34.5 | 19.4 – 33.1 |
|  | mean (SD) | 25.1 (4.1) | 24.9 (3.0) |
|  | BMI > 25 | 11 | 10 |
|  | BMI > 30 | 2 | 2 |
| **Waist circumference (in)** | Waist circumference, mean (SD) | 36.1 (5.0) | 35.4 (4.5) |
| **Systolic BP (mmHg)** | range | 99.0 – 170.2 | 98.5 – 168.2 |
|  | mean (SD) | 131.1 (20.1) | 125.7 (16.6) |
| **Diastolic BP (mmHg)** | range | 65.0 – 116.5 | 66.8 – 104.2 |
|  | mean (SD) | 83.2 (12.1) | 81.6 (9.2) |
| **Heart rate (bpm)** | range | 52.5 – 84.2 | 53.2 – 91.0 |
|  | mean (SD) | 70.2 (9.3) | 70.1 (10.6) |
|  | Smoking (metabolite-based) | 0 | 2 |
| **Baseline NAD+** | Baseline icNAD (uM) | 28.8 (4.7) | 28.9 (4.8) |
|  | Baseline cirNAD (nM) | 390.2 (94.5) | 392.8 (105.9) |
|  | Vitamin D supplementation | 14 (60.9%) | 8 (29.6%) |

Values are mean (SD) for continuous variables and n (%) for categorical variables. Continuous: age, BMI, weight, height, systolic BP, diastolic BP, heart rate. Categorical: sex, race (White, Asian, Black or African American, Multiracial), ethnicity. Baseline icNAD (uM) and cirNAD (nM) values averaged across the two pre-treatment blood draws. Groups were well balanced at randomization. Primary analysis cohort (N = 50).

**Table 2.** Primary endpoint NAD concentrations by treatment group and timepoint.

| Analyte | Visit | N (LNAD+) | LNAD+ Mean (SD) | N (Placebo) | Placebo Mean (SD) |
| --- | --- | --- | --- | --- | --- |
| cirNAD (nM) | Baseline | 23 | 389.9 (86.2) | 27 | 391.4 (105.1) |
|  | Day 4 | 23 | 390.3 (100.2) | 27 | 378.5 (114.5) |
|  | Day 6 | 22 | 386.5 (103.2) | 26 | 408.0 (119.5) |
| icNAD (μM) | Baseline | 23 | 28.7 (4.6) | 27 | 29.0 (4.7) |
|  | Day 4 | 23 | 39.1 (5.5) | 27 | 28.4 (4.8) |
|  | Day 6 | 22 | 43.5 (7.1) | 26 | 28.2 (4.7) |

Descriptive statistics (mean, SD) for intracellular NAD (icNAD, μM) and circulating NAD (cirNAD, nM) at Baseline, Day 4, and Day 6 by treatment group. MMRM estimates presented in manuscript text.

### Primary Endpoints: Intracellular and Circulating NAD+

Participants were compliant with study procedures. Manual verification of paper and/or electronic dose logs confirmed treatment compliance for 49 out of 51 participants (96%). The study met its primary endpoint: oral LNAD+ administration resulted in a statistically significant increase in intracellular NAD (icNAD) concentrations relative to placebo across both on-protocol visits (Day 4: MMRM estimate 38.2%, 95% CI: 29.1–48.0%, p = 4.11 × 10^−12^; Day 6: MMRM estimate 53.0%, 95% CI: 41.2–65.7%, p = 5.48 × 10^−14^; Hedges’ g = 3.66).

At Day 6, mean icNAD in the LNAD+ group was 43.5 μM (SD 7.1) compared to 28.2 μM (SD 4.7) in the placebo group, whose levels remained unchanged from baseline (29.0 μM, SD 4.7). A total of 18 out of 23 participants (78.3%) receiving LNAD+ achieved greater than 30% increase in icNAD by Day 6, with a minimum individual increase of 9.4% (**Figure 1**). The treatment effect was confirmed by nonparametric BCa bootstrap (bootstrap 95% CI: 11.5–18.0 μM raw difference; 1,000,000 BCa iterations).

**Figure 1.**
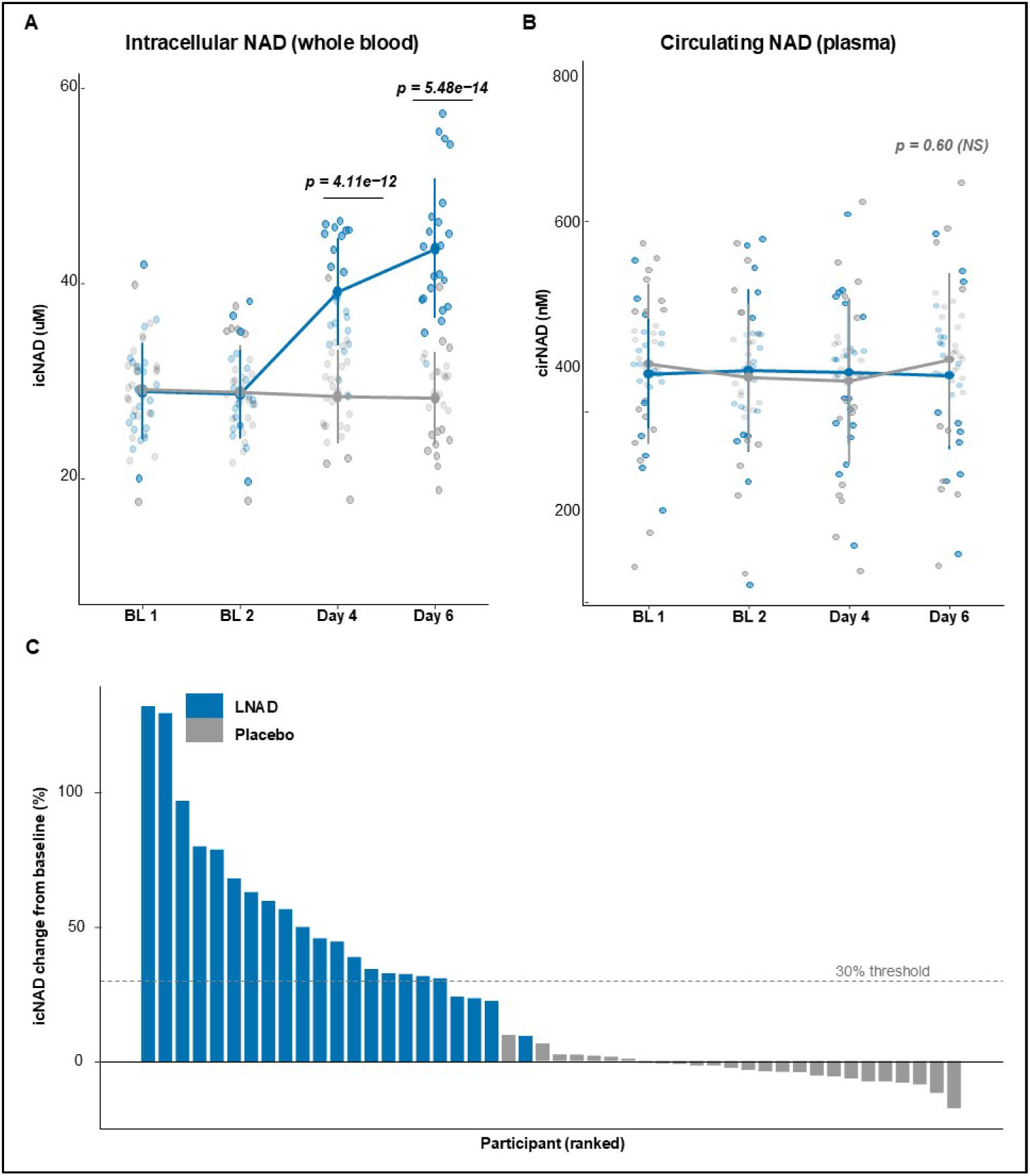
Study design and primary NAD+ endpoints. (A) Intracellular NAD (icNAD, μM) measured in whole blood at Baseline 1, Baseline 2, Day 4, and Day 6 in LNAD+ (blue, n = 23) and Placebo (gray, n = 27) groups. Individual data points are shown with group means ± SD connected by lines. MMRM treatment contrast p-values annotated at Day 4 and Day 6. (B) Circulating NAD (cirNAD, nM) measured in plasma. (C) Waterfall plot of individual icNAD percent change from baseline at Day 6, ranked by magnitude. Blue bars = LNAD+; gray bars = Placebo. Dashed line indicates 30% responder threshold. Primary analysis cohort (N = 50).

LNAD+ administration did not significantly affect circulating NAD (cirNAD) at either timepoint (Day 4: p = 0.43; Day 6: p = 0.60), with plasma NAD concentrations remaining stable in both groups throughout the study period. Full model results for icNAD and cirNAD are presented in **Supplementary Tables 4-5**. Bootstrap distributions of treatment effects are presented in **Supplementary Figures 3-6**.

### Secondary Endpoint: NAD+ Metabolites

LNAD+ administration led to differential, statistically significant increases in plasma abundance of 1-methylnicotinamide (MeNAM; Day 4: p = 6.29 × 10^−17^, 551% estimated group difference; Day 6: p = 5.39 × 10^−13^, 364%) and N1-methyl-2-pyridone-5-carboxamide (2PY; Day 4: p = 2.55 × 10^−19^, 512%; Day 6: p = 2.95 × 10^−16^, 368%), compared to placebo (**Figure 2**). Nicotinamide (NAM) showed a significant treatment effect at Day 4 (p = 0.016, 54% estimated group difference) that was not maintained at Day 6 (p = 0.30). No significant changes were detected in the plasma abundance of quinolinate or trigonelline, metabolites associated with the de novo and Preiss–Handler NAD+ biosynthetic pathways, respectively. MeNAM and 2PY remained significant after corrections for multiple testing. Pre- and post-intervention levels of NAD+ metabolites are reported in **Supplementary Table 6**.

**Figure 2.**
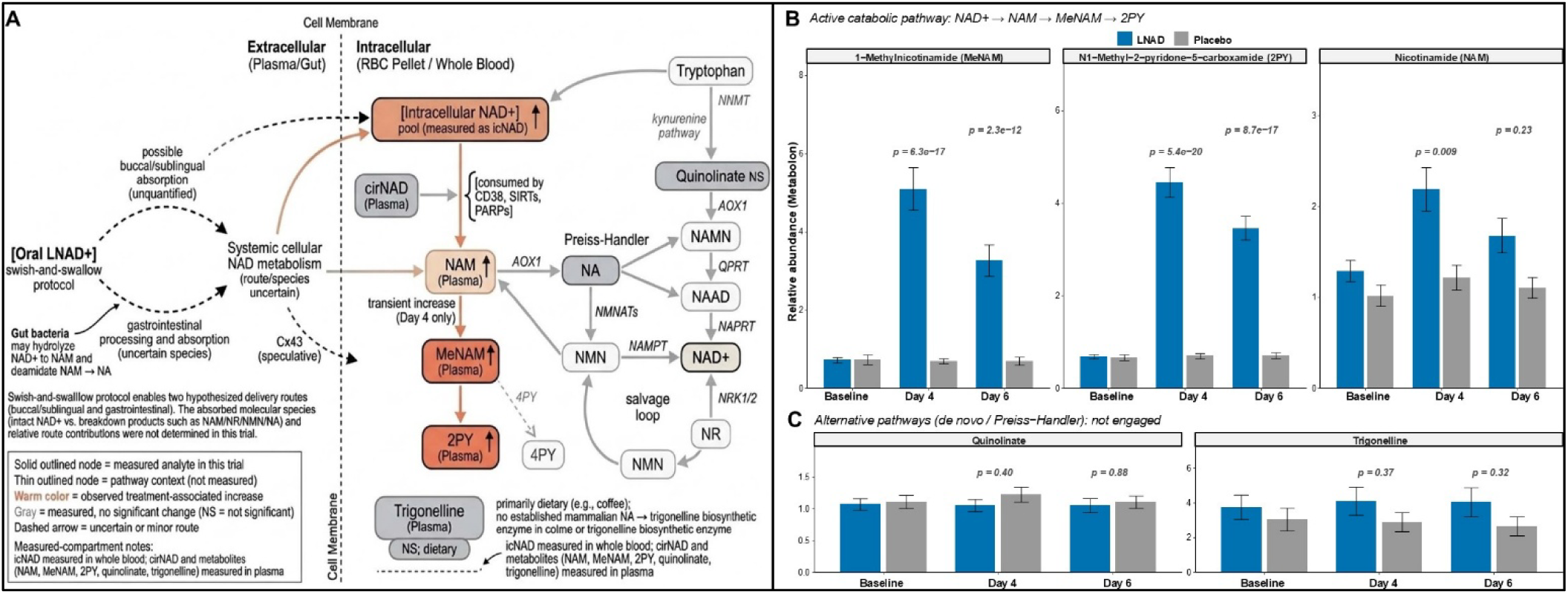
NAD+ metabolic flux: metabolite pattern consistent with enhanced catabolic processing. (A) Interpretive pathway schematic showing NAD+ biosynthetic and catabolic routes with measured analytes highlighted. Solid-outlined nodes indicate metabolites quantified in this trial; thin-outlined nodes are contextual pathway intermediates (not measured). Warm colors indicate observed treatment-associated increases; gray indicates measured but unchanged. Dashed arrows indicate uncertain or minor routes. This swish-and-swallow protocol may permit both oral mucosal exposure and gastrointestinal delivery; however, relative contributions were not determined in this trial; (B) Active catabolic pathway metabolites: 1-methylnicotinamide (MeNAM), N1-methyl-2-pyridone-5-carboxamide (2PY), and nicotinamide (NAM). Grouped bar charts showing Metabolon median-normalized relative abundance (mean ± SEM) at Baseline, Day 4, and Day 6 by treatment group (LNAD+ blue, Placebo gray). MMRM p-values annotated. (C) Quinolinate (de novo synthesis intermediate) and trigonelline (primarily dietary). Primary analysis cohort (N = 50).

### Secondary Endpoints: Clinical Laboratory Tests

Total serum bilirubin demonstrated a nominally significant decrease in the LNAD+ group relative to placebo at Day 6 (MMRM p = 0.019, g = −0.80). Albumin also showed a nominally significant decrease (p = 0.033, g = −0.77), as did the albumin/globulin (A/G) ratio (p = 0.045, g = −0.68), with a borderline trend for alkaline phosphatase (p = 0.067, g = −0.63). These correlated hepatic markers all remained within normal clinical reference ranges, and liver transaminases were unchanged (ALT: p = 0.80; AST: p = 0.29; GGT: p = 0.86), providing no evidence of hepatocellular injury. These findings did not survive multiplicity correction (all adjusted p > 0.31).

In the hematological panel, mean corpuscular hemoglobin (MCH; p = 0.031, g = −0.72) and mean corpuscular hemoglobin concentration (MCHC; p = 0.028, g = −0.73) both registered nominally significant reductions compared to placebo, at Day 6. These shifts, while statistically detectable, were small in absolute magnitude and remained within normal clinical ranges.

LNAD+ administration was associated with a nominally significant increase in absolute lymphocyte count at Day 6 (MMRM estimate = 0.125 × 10^9^/L, Hedges’ g = 0.73, p = 0.037), with a trend in lymphocyte percentage (+4.36 percentage points, g = 0.69, p = 0.053). This effect was not present at Day 4 (absolute lymphocytes: p = 0.58; percentage: p = 0.33). The lymphocyte increase was not accompanied by detectable inflammatory activation. Tumor necrosis factor-α (TNF-α) was unchanged (p = 0.127, g = −0.49), as were interleukin-6 (IL-6; p = 0.866, g = −0.07), high-sensitivity C-reactive protein (hs-CRP; p = 0.913, g = 0.04), and erythrocyte sedimentation rate (ESR; p = 0.412, g = −0.28). Total white blood cell count was stable (p = 0.687), and neutrophil percentage showed a nonsignificant downward trend (g = −0.59, p = 0.093), while neutrophil absolute counts were unchanged (p = 0.334). The basophil percentage was nominally elevated (g = 0.80, p = 0.016), though this was not confirmed in absolute counts (p = 0.329), consistent with a proportional shift secondary to differential changes rather than a true basophil response. The lymphocyte finding did not survive Benjamini–Hochberg correction within the laboratory outcome family (adjusted p = 0.317).

Finally, with respect to oxidative stress, no treatment-related changes were observed in serum gamma-glutamyl transferase (GGT; Day 6 p = 0.86) or cumulative levels of reactive oxygen metabolites (ROM; p = 0.94).

No other clinical laboratory tests displayed nominal statistical significance at Day 6. Renal function (creatinine: p = 0.14; eGFR: p = 0.25; BUN: p = 0.83), lipid panels (total cholesterol: p = 0.65; LDL: p = 0.29; HDL: p = 0.67; triglycerides: p = 0.70), electrolytes, and glucose metabolism markers were all unaffected. No nominally significant laboratory findings survived Benjamini–Hochberg correction within their respective outcome families (all adjusted p > 0.31). Pre- and post-intervention levels of clinical laboratory panels are reported in **Supplementary Table 7**. Complete Day 4 and Day 6 treatment contrasts across all endpoints are presented in **Figure 3**.

**Figure 3.**
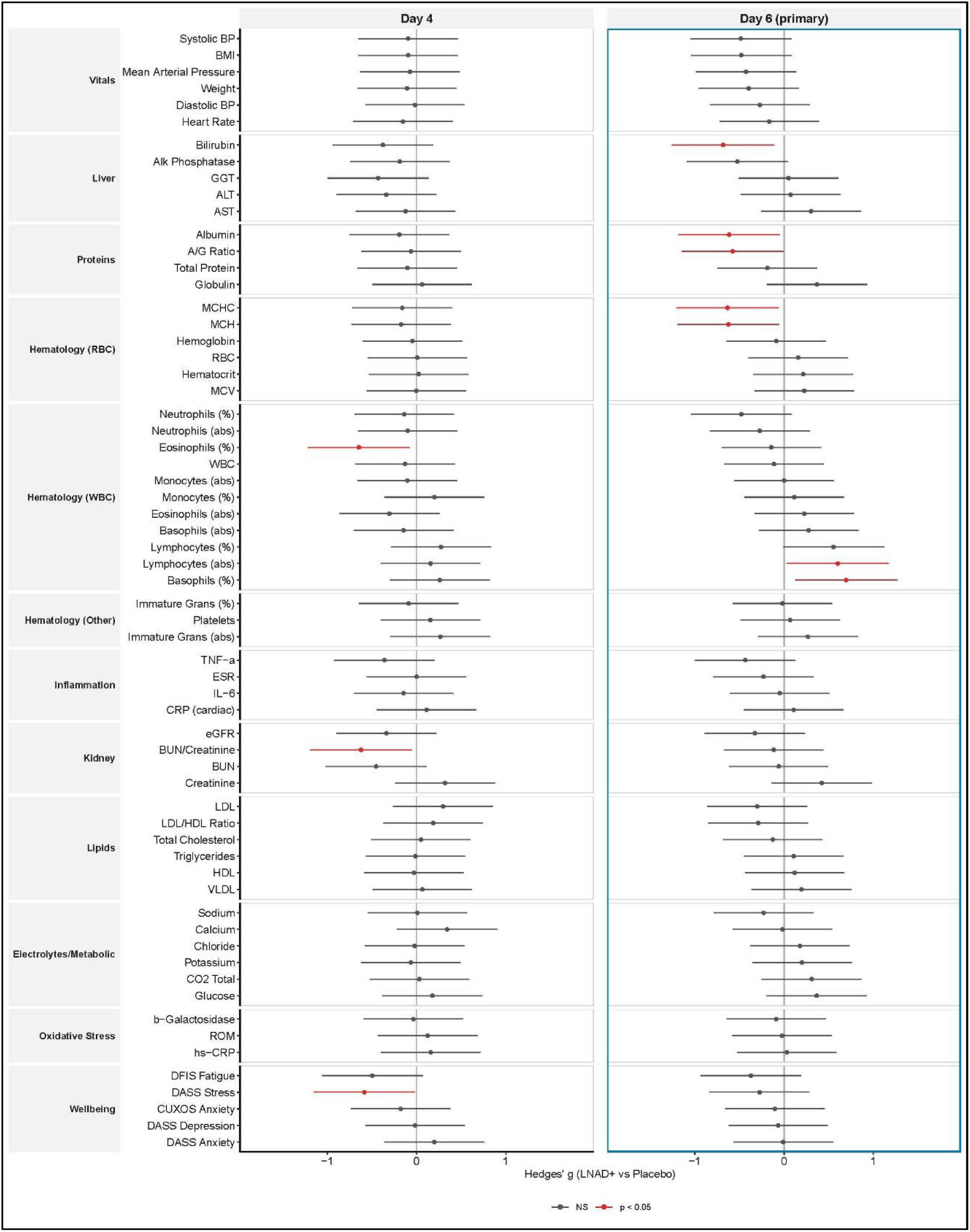
Safety forest plot: Day 4 and Day 6 MMRM treatment contrasts across endpoints. Hedges’ g effect size estimates (LNAD+ vs Placebo) with 95% confidence intervals for clinical laboratory, vital sign, and wellbeing endpoints, presented as side-by-side panels: Day 4 (left) and Day 6 (right). Hedges’ g computed with small-sample correction factor J = 1 − 3/[4(n□ + n□ − 2) − 1]; CIs from large-sample variance approximation SE(g) ≈ √(1/n□ + 1/n□ + g²/[2(n□ + n□)]). Endpoints organized by clinical domain: Vitals, Liver, Proteins, Hematology (RBC, WBC, Other), Inflammation, Kidney, Lipids, Electrolytes/Metabolic, Oxidative Stress, Wellbeing. Red points indicate nominal p < 0.05; gray points indicate p ≥ 0.05. No secondary findings survived Benjamini–Hochberg correction within their respective outcome families. Vertical line at zero represents no treatment effect. Primary analysis cohort (N = 50).

### Secondary Endpoint: Safety and Tolerability

Systolic blood pressure (SBP) showed a nonsignificant trend toward reduction in the LNAD+ group at Day 6 (MMRM estimate −7.1 mmHg, g = −0.61, p = 0.092), with no significant changes in diastolic blood pressure (p = 0.34) or heart rate (p = 0.55). Body weight and BMI showed nonsignificant decreasing trends (BMI: p = 0.092, g = −0.58; weight: p = 0.163, g = −0.48).

No differential changes were observed in measures of subjective well-being (DASS-21 Depression: p = 0.82; DASS-21 Anxiety: p = 0.97; DFIS Fatigue: p = 0.19; CUXOS Anxiety: p = 0.71). DASS-21 Stress showed a nominally significant improvement at Day 4 (p = 0.0435) that did not persist to Day 6 (p = 0.468). Wearable-derived activity endpoints showed nominal shifts toward higher activity and lower sedentary time (e.g., very active minutes increased and sedentary minutes decreased at Day 6), but no wearable endpoint remained significant after hierarchical false discovery rate correction; sleep metrics were unchanged. Full results of wearable endpoints are presented in **Supplementary Table 8**.

Only one treatment-emergent adverse event was recorded: mild nausea (Grade 1, CTCAE v5.0) in a single participant in the LNAD+ arm on Day 2, resolving within 24 hours after discontinuation. The overall on-protocol symptom burden, assessed via a comprehensive 71-item Review of Systems questionnaire, was comparable between groups (LNAD+ 2.92% vs Placebo 2.40% symptom-day incidence; GLMM p = 0.68). **Table 3** and **Table 4** present key descriptives for adverse events and symptom self-reports, and clinical / vital sign endpoints with nominal p < 0.10 at Day 6, respectively. A detailed summary of ROS symptom reports is presented in **Supplementary Table 9**. Full results of ROS GLMM analyses are presented in **Supplementary Table 10** and **Supplementary Figure 7**.

**Table 3.**
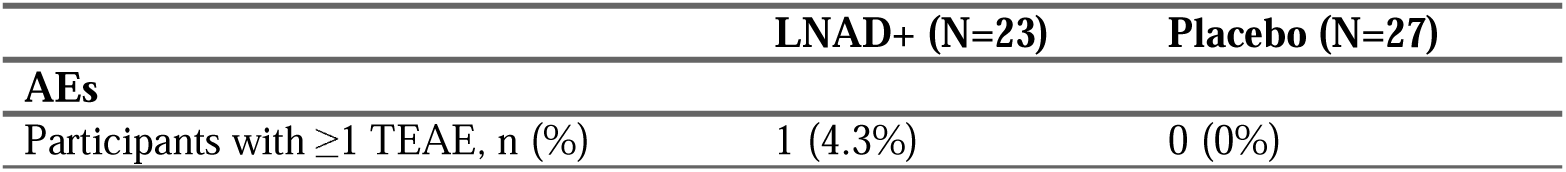
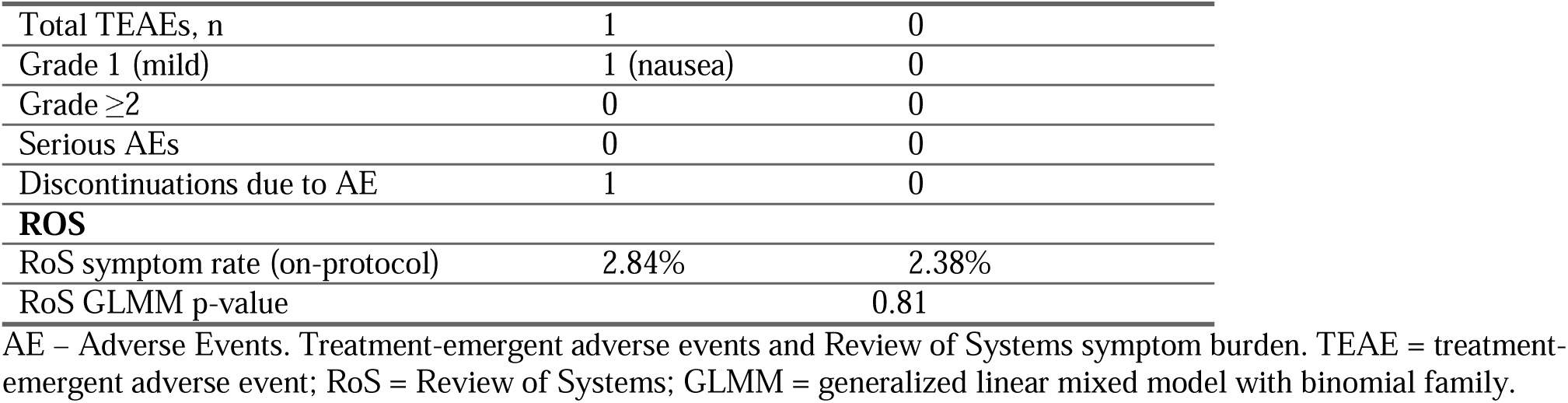
Treatment-emergent adverse events and Review of Systems symptom burden.

**Table 4.** Clinical laboratory and vital sign endpoints with nominal p < 0.10 at Day 6.

| Domain | Analyte | LNAD+ Day 6 | Placebo Day 6 | Hedges' g | MMRM p | Direction |
| --- | --- | --- | --- | --- | --- | --- |
| Hematology (WBC) | Basophils (%) | 1.05 (0.49) | 0.80 (0.50) | 0.80 | 0.0164 | ↑ LNAD+ |
| Liver | Bilirubin, Total | 0.36 (0.18) | 0.46 (0.39) | -0.80 | 0.0188 | ↓ LNAD+ |
| Hematology (RBC) | MCHC | 33.74 (0.78) | 34.07 (0.76) | -0.73 | 0.0283 | ↓ LNAD+ |
| Hematology (RBC) | MCH | 30.44 (1.28) | 30.44 (1.48) | -0.72 | 0.0307 | ↓ LNAD+ |
| Proteins | Albumin | 4.45 (0.32) | 4.52 (0.28) | -0.77 | 0.0326 | ↓ LNAD+ |
| Hematology (WBC) | Lymphocytes (abs) | 1.89 (0.50) | 1.56 (0.47) | 0.73 | 0.0366 | ↑ LNAD+ |
| Proteins | A/G Ratio | 1.91 (0.32) | 2.02 (0.31) | -0.68 | 0.0448 | ↓ LNAD+ |
| Hematology (WBC) | Lymphocytes (%) | 35.00 (7.26) | 29.32 (7.30) | 0.69 | 0.0532 | ↑ LNAD+ |
| Liver | Alk Phosphatase | 71.18 (19.98) | 75.20 (19.58) | -0.63 | 0.0674 | ↓ LNAD+ |
| Vitals | Systolic BP | 125.6 (16.1) | 126.3 (19.4) | -0.61 | 0.0921 | ↓ LNAD+ |
| Vitals | BMI | 25.2 (4.0) | 25.0 (3.1) | -0.58 | 0.0924 | ↓ LNAD+ |
| Hematology (WBC) | Neutrophils (%) | 52.32 (8.93) | 58.80 (8.76) | -0.59 | 0.0927 | ↓ LNAD+ |

Note. Hedges' g computed from MMRM contrasts. No findings survived mEff correction within their respective outcome families. All values remained within clinical reference ranges.

### Exploratory Endpoint: Baseline NAD+ Correlation Signatures

Exploratory Spearman rank correlations between baseline NAD+ concentrations and clinical endpoints tentatively supported compartment-specific signatures that reinforce the rationale for separating intracellular and circulating NAD+ measurements (**Supplementary Table 11**). Baseline cirNAD was correlated with tumor necrosis factor-α (ρ = −0.56, p_BH_ = 0.003), creatinine (ρ = −0.51, p_BH_ = 0.006), and positively correlated with HDL cholesterol (ρ = 0.43, p_BH_ = 0.042), forming a metabolic and anti-inflammatory signature consistent with CD38-mediated NAD+ consumption in inflammatory states. Baseline icNAD, by contrast, tracked red blood cell indices - hemoglobin (ρ = 0.54, p_BH_ = 0.003) and hematocrit (ρ = 0.50, p_BH_ = 0.006) - reflecting the measurement compartment (whole blood), and with the salvage pathway substrate nicotinamide (NAM; ρ = 0.47, p_BH_ = 0.018), suggesting that individuals with higher circulating nicotinamide availability maintain higher intracellular NAD+ pools.

## DISCUSSION

This double-blind, randomized, placebo-controlled Phase 0/1b trial demonstrated that 5 days of oral LathMized® NAD+ (LNAD+) produce a rapid, substantial increase in intracellular NAD (icNAD) measured in whole blood, without a corresponding increase in plasma NAD. In parallel, large increases in the downstream catabolites 1-methylnicotinamide (MeNAM) and N1-methyl-2-pyridone-5-carboxamide (2PY) provide pharmacodynamic evidence that administered NAD engages active metabolic processing pathways: both remained significant after false discovery rate correction. Across comprehensive clinical laboratory panels, vital signs, patient-reported outcomes, and wearable-derived activity and sleep measures, LNAD+ was well tolerated over the short dosing period. Secondary clinical signals observed are reported transparently and should be interpreted as hypothesis-generating.

### Cellular Uptake and Absorption Biology

A key question raised by these findings is how exogenous oral NAD+ reaches the intracellular compartment. We outline several non-exclusive mechanisms that may contribute. First, extracellular NAD+ can be hydrolyzed by ectoenzymes (CD38, CD73, CD157) to yield nicotinamide (NAM), nicotinamide riboside (NR), or nicotinamide mononucleotide (NMN), each of which has established cellular uptake pathways (46, 47). Second, intact NAD+ may cross cell membranes via connexin 43 hemichannels under calcium-regulated conditions (48), though the quantitative contribution of this pathway *in vivo* at physiological concentrations remains uncertain. Third, recent evidence from head-to-head precursor trials demonstrates that oral NAD+ precursors undergo extensive gut microbial conversion (NMN and NR are deamidated to nicotinic acid derivatives before systemic absorption (22, 49), suggesting that the route from oral NAD+ to intracellular NAD+ may involve multiple intermediate conversions rather than direct intact-molecule uptake. Thus, a plausible alternative explanation is that orally administered NAD+ is substantially hydrolyzed prior to absorption and functions primarily as a NAM source. Importantly, recent human data show that oral NAD boosters can enter host NAD metabolism via distinct, microbiome-dependent routes (e.g., deamidation to nicotinic acid derivatives), underscoring that “oral NAD booster” to “NAM-only equivalence” is not a safe assumption (22, 49).

The present study cannot determine contributions among these mechanisms as we did not measure plasma NR, NMN, or nicotinic acid (NA) concentrations, gut microbial conversion, or erythrocyte transporter expression. However, the empirical finding that the intracellular pool rises while the plasma pool does not, and that downstream catabolites accumulate, is consistent with efficient cellular NAD augmentation rather than simple extracellular persistence.

### Intracellular NAD+ Elevation and Metabolic Flux

NAD+ metabolism is increasingly conceptualized in terms of dynamic flux rather than static pool sizes (50). NAD flux represents the dynamic turnover of NAD through biosynthetic, salvage, and consumption pathways mediated by enzymes such as sirtuins, PARPs, and CD38. Because these pathways operate within metabolically active cells, intracellular NAD measurements may better reflect NAD metabolic flux than whole-blood measurements.

The approximately 53% increase in whole blood (primarily RBC-driven) NAD+ observed at Day 6 is commensurate with effects reported for NAD+ precursor interventions of longer duration, where whole-blood NAD+ increases of 22–100% with nicotinamide riboside (NR) and 11–60% with nicotinamide mononucleotide (NMN) are typically observed following 2–12 weeks of treatment (25, 49, 51–53). Moreover, the magnitude and rapidity of the icNAD response - detectable at Day 4 (38.2% group difference) and increasing by Day 6 - suggests efficient delivery of intact or minimally processed NAD+ to the cellular compartment. Considering a large effect threshold of g>0.80 (45), the effect size observed (Hedge’s g = 3.66) is exceptionally large and indicates virtually no overlap between the treatment and placebo distributions at Day 6.

The null effect on circulating NAD (cirNAD) is informative as well, as exogenous NAD+ is not accumulating in the extracellular space but is being taken up by cells or undergoing extracellular degradation with subsequent cellular uptake of degradation products. This compartment-selective profile contrasts with intravenous NAD, which elevates circulating NAD transiently. The substantial elevation of NAD+ catabolites MeNAM and 2PY provides further preliminary pharmacodynamic evidence that exogenous NAD+ enters active metabolic processing.

### Safety and Clinical Implications

LNAD+ demonstrated a favorable safety profile across all assessed domains. The single treatment-emergent adverse event—mild nausea (Grade 1, CTCAE v5.0)—resolved within 24 hours and is consistent with the known gastrointestinal tolerability profile of oral PEG-containing or high-dose oral formulations (37). The overall on-protocol symptom burden, assessed via a 71-item Review of Systems questionnaire, was comparable between groups (GLMM p = 0.81), providing no evidence of systematic symptom induction.

The nominally significant increase in absolute lymphocyte count (g = 0.73, p = 0.037) did not survive multiplicity correction. Even so, the observation that the lymphocyte increase occurred without concurrent elevation of inflammatory markers TNF-α & CRP is notable, because an inflammatory etiology (infection, autoimmunity, or nonspecific immune activation) would typically result in concurrent cytokine and acute-phase reactant elevation. One possible interpretation is that restored NAD+ availability supports lymphocyte metabolic fitness (viability, homeostatic proliferation, or resistance to apoptosis) independently of inflammatory signaling. This interpretation is supported by recent evidence that NAD+ depletion in aged CD8+ T cells drives mitochondrial dysfunction and loss of stem-like properties, and that NAD+ precursor supplementation rescues these phenotypes without activating inflammatory pathways (13, 54). The concurrent downward trend in neutrophil percentage (g = −0.59, p = 0.093) is compatible with this framing, as a shift in the lymphocyte-to-neutrophil ratio is associated with reduced systemic inflammatory tone. Separately, nominally significant shifts in MCH (p = 0.031) and MCHC (p = 0.028) - RBC-intrinsic indices not influenced by plasma volume - were driven primarily by placebo-arm increases rather than LNAD+ decreases; these small changes of unclear clinical significance remained within reference ranges.

Among clinical laboratory parameters, nominally significant decreases in albumin (p = 0.033, g = −0.77), A/G ratio (p = 0.045, g = −0.68), and total bilirubin (p = 0.019, g = −0.80) were observed in the LNAD+ group relative to placebo. None survived multiplicity correction. For protein markers (albumin, A/G ratio), arm-specific trajectories show bidirectional divergence at Day 6: the LNAD+ arm decreases mildly from baseline while the placebo arm drifts upward. Total bilirubin showed the largest nominal effect among these markers (g = −0.80, p = 0.019). Unlike albumin, bilirubin is a heme degradation catabolite whose clearance depends on hepatic glucuronidation via UGT1A1, the major bilirubin-glucuronidation enzyme. One hypothesis is that enhanced intracellular NAD+ availability upregulates UGT1A1 cofactor supply (UDP-glucuronic acid is synthesized through NAD+-dependent oxidation of UDP-glucose) and/or UGT1A1 expression via the SIRT1–PGC-1α axis, accelerating bilirubin conjugation. This would represent a pharmacodynamic consequence of NAD+ repletion.

However, this study did not report on conjugated vs. unconjugated bilirubin fractions, UGT1A1 activity, or hepatic NAD+ status, and the bilirubin decrease could also reflect plasma volume changes, analytic variability, or chance. Importantly, hepatocellular injury markers were entirely unchanged (ALT: p = 0.80; AST: p = 0.29; GGT: p = 0.86), and all observed values remained within clinical reference ranges, providing no evidence of hepatotoxicity. These patterns should be monitored in longer-duration studies with pre-specified endpoints, including bilirubin fractionation, plasma volume markers, and UGT1A1 genotype/activity to distinguish among these interpretations.

The elevation of MeNAM and 2PY has implications for specific clinical populations. Both metabolites have been previously identified as potential uremic toxins that accumulate in chronic kidney disease (CKD), although their role in cardiovascular and renal pathology is under investigation (55).

While the observed increases are in healthy adults with presumably normal renal clearance, future studies of NAD+ supplementation in renally impaired populations should include systematic monitoring of these catabolites.

### Limitations

Several limitations should be considered when interpreting these findings. The sample size (n = 50 primary analysis; n = 51 full ITT sensitivity set) was sufficient to detect the very large primary endpoint effect on icNAD but was not powered for smaller secondary effects. Multiple comparisons also constrain inference for secondary outcomes: across the 70 clinical laboratory, vital sign, and wellbeing endpoints evaluated in the primary analysis pipeline, the effective number of independent tests was M_eff_ = 56 (Galwey eigenvalue method with Ledoit–Wolf shrinkage), corresponding to a global Šidák threshold of α ≈ 0.000926. Under this global correction, only icNAD and NAD metabolites MeNAM and 2PY remained statistically significant. Other nominal findings within this endpoint set (e.g., lymphocytes, hepatic marker shifts, SBP trend) should be interpreted as hypothesis-generating and prioritized for replication.

We acknowledge that the 5-day dosing period establishes acute pharmacodynamic proof-of-concept but cannot address questions of chronic NAD+ homeostasis, potential compensatory downregulation of endogenous NAD+ biosynthesis, or sustained clinical benefit. The study population (predominantly White, healthy adults aged 45–75 from a single site in the Pacific Northwest of the United States) also limits its generalizability. The absence of severely metabolically compromised participants (e.g., with type 2 diabetes, sarcopenia, or neurodegenerative disease) means the study does not directly translate to the populations most likely to benefit from NAD+ restoration, where baseline NAD+ depletion may produce larger treatment effects.

The Jinfiniti assay measures total NAD (NAD+ + NADH) rather than the NAD+/NADH ratio. If NADH is co-elevated, the functional redox implications would differ from a selective increase in oxidized NAD+. Future studies should incorporate ratio-specific assays or complementary measures of NAD+-dependent enzymatic activity to resolve this question.

The analytical approach evolved from the original SAP in five documented respects (see Methods). The most substantive change (adoption of MMRM over mixed ANOVA) represents a more conservative analytical framework that is now standard in longitudinal clinical trial analysis. The shift from one-tailed to two-tailed testing further increases inferential stringency. All primary findings are robust across the original and updated analytical frameworks, across the full (N = 51) and sensitivity (N = 50) analysis sets, and across parametric (MMRM) and nonparametric (BCa bootstrap) inference methods.

Finally, we recognize that measurement of icNAD in the whole blood (dominated by RBCs rather than PBMCs) has both advantages and limitations for interpreting the biological significance of the observed increase. Mature erythrocytes have limited NAD+ biosynthetic capacity: they lack nuclei and mitochondria and therefore cannot perform de novo NAD+ synthesis, relying exclusively on the NAMPT-NMN-NMNAT3 salvage axis, as demonstrated by studies showing that NMNAT3 deficiency depletes RBC NAD+ and causes hemolytic anemia (56). Erythrocyte NAD+ concentrations decline with advancing age in humans (57). Consequently, elevation of RBC NAD+ following oral administration implies effective delivery of NAD+ (or its immediate metabolites) across the erythrocyte membrane, a finding with implications for other cell types with similarly constrained biosynthetic capacity. We acknowledge that generalization of the signal from erythrocytes to metabolically active tissues such as immune cells, muscle, liver, or brain requires tissue-specific measurements that were beyond the scope of this study.

## Conclusion

RENEWAL-NAD+ was a Phase 0/1b randomized, double-blind, placebo-controlled trial demonstrating that 5 days of oral LathMized® NAD+ (LNAD+) produces a rapid, substantial increase in intracellular NAD (icNAD) measured in the whole blood compartment, while circulating NAD (cirNAD) in plasma remains unchanged. The marked increases in the downstream NAD catabolites 1-methylnicotinamide (MeNAM) and N1-methyl-2-pyridone-5-carboxamide (2PY) provide pharmacodynamic evidence of engagement of NAD metabolic flux pathways following oral dosing.

LNAD+ was well tolerated over the short intervention, with no clinically meaningful safety concerns observed across regular and comprehensive assessment of clinical laboratory tests, vitals, and subjective indices of well-being. Secondary signals (including a nominal Day 6 lymphocyte increase without inflammatory marker elevation) did not remain significant after multiplicity correction across the 70 clinical laboratory, vital sign, and wellbeing endpoints analyzed in the primary pipeline and should be interpreted as hypothesis-generating. Additional exploratory multi-omic profiling of pharmacodynamic responses to LNAD+ are ongoing and will be reported separately. Together, the findings establish short-term human intracellular NAD augmentation and downstream metabolic engagement after oral NAD+ administration, supporting longer-duration studies with dose-response characterization, tissue- and cell-specific NAD measurements, and pre-specified immune and hepatic monitoring endpoints.

## ABBREVIATIONS

2PY: N1-methyl-2-pyridone-5-carboxamide
ALP: Alkaline Phosphatase
ALT: Alanine Transaminase
AST: Aspartate Transaminase
BCa: Bias-Corrected and Accelerated
BMI: Body Mass Index
BUN: Blood Urea Nitrogen
CD38: Cluster of Differentiation 38
CI: Confidence Interval
cirNAD: Circulating/Extracellular NAD
CLIA: Clinical Laboratory Improvement Amendments
CTCAE: Common Terminology Criteria for Adverse Events
CUXOS-D: Clinically Useful Anxiety Outcome Scale, Daily Version
CV: Coefficient of Variation
D-FIS: Daily Fatigue Impact Scale
DASS-21: Depression, Anxiety, and Stress Scale-21
DBP: Diastolic Blood Pressure
eGFR: Estimated Glomerular Filtration Rate
ESR: Erythrocyte Sedimentation Rate
GGT: Gamma-Glutamyl Transferase
GLMM: Generalized Linear Mixed Model
HDL: High-Density Lipoprotein
hs-CRP: High-Sensitivity C-Reactive Protein
icNAD: Intracellular NAD
IL-6: Interleukin-6
LDL: Low-Density Lipoprotein
LNAD+: LathMized® NAD+
MAR: Missing at Random
MCH: Mean Corpuscular Hemoglobin
MCHC: Mean Corpuscular Hemoglobin Concentration
MeNAM: 1-Methylnicotinamide
M_eff_: Effective Number of Independent Tests
MMRM: Mixed Models for Repeated Measures
NAD+: Nicotinamide Adenine Dinucleotide
NAM: Nicotinamide
NMN: Nicotinamide Mononucleotide
NNMT: Nicotinamide N-Methyltransferase
NR: Nicotinamide Riboside
PARP: Poly(ADP-ribose) Polymerase
PEG: Polyethylene Glycol
QID: Quarter-in-die (four times daily)
RBC: Red Blood Cell
ROM: Reactive Oxygen Metabolites
RoS: Review of Systems
SAP: Statistical Analysis Plan
SBP: Systolic Blood Pressure
SIRT: Sirtuin
TNF-α: Tumor Necrosis Factor Alpha
UGT1A1: UDP-Glucuronosyltransferase 1A1
US: Unstructured (covariance)
WBC: White Blood Cell Count

## AUTHOR CONTRIBUTIONS

*Study Design*: Sergey A. Kornilov, Andrew Magis, Wendy Komac; *Study Conception*: Wendy Komac; *Sample Acquisition*: Steve M. Coppess, Andrew Magis, & Sergey A. Kornilov; *Statistical Analysis*: Sergey A. Kornilov & Waylon J. Hastings; *Data Interpretation and Manuscript Drafting*: Waylon J. Hastings, Sergey A. Kornilov, Michael Leitz-Langan, Lynne Fahey McGrath; *Manuscript Critical Review*: All authors.

## ACKNOWLEDGEMENTS

We thank the staff at Institute for Systems Biology and ISB BioAnalytica Inc. for their diligence in conducting participant screening, recruitment, and study procedures as well as all research participants who provided their time and resources to complete RENEWAL-NAD+. We wish to extend our sincerest appreciation to Dr. Mia Nease for her generous contribution of time, expertise, and scientific insight in guiding the incorporation of multi-omics approaches into the RENEWAL Study design. Her thoughtful input and collaborative spirit were invaluable to the development of this work.

## ETHICAL STATEMENT

This study, and any subsequent modifications, was reviewed and approved by the Western Institutional Review Board (now known as WCG-IRB), as defined by Federal Regulatory Guidelines (Ref. Federal Register Vol. 46, No. 17, January 27, 1981, part 56) and the Office for Protection from Research Risks Reports: Protection of Human Subjects (Code of Federal Regulations 45 CFR 46). IRB Tracking Number 20215675. The study was conducted per IRB-approved protocol (10/13/2021, Version 1) with study code ISBA-101 and study number 1320748. The study was conducted according to the ethical principles of the Declaration of Helsinki and current Good Clinical Practice guidelines, and in compliance with federal, state, and local regulatory requirements, including 21 CFR 312, 21 CFR 50, and 21 CFR 56.

## CONSENT

Informed consent was obtained from each participant in an electronic format (e-consent) at enrollment following pre-screening and pending confirmation with the Study Coordinator. The Study Coordinator and/or the PI of the study described the study and answered questions about the study goals and the protocol, as needed. The principles of informed consent followed are described by Federal Regulatory Guidelines (Federal Register Vol. 46, No. 17, January 27, 1981, part 50) and the Office for Protection from Research Risks Reports: Protection of Human Subjects (Code of Federal Regulations 45 CFR 46). The Informed Consent Form was reviewed and approved by Western IRB (now known as WCG IRB) as part of the protocol submission.

## DATA AVAILABILITY

The clinical trial data underlying this study are available from the corresponding author upon reasonable request, subject to participant privacy protections and institutional data sharing agreements. The reproducible analysis pipeline code is available at [repository URL to be added prior to publication].

## REFERENCES

1. Belenky P, Bogan KL, Brenner C. NAD+ metabolism in health and disease. Trends Biochem Sci. 2007;32(1):12–9.

2. Verdin E. NAD(+) in aging, metabolism, and neurodegeneration. Science. 2015;350(6265):1208–13.

3. Canto C, Menzies KJ, Auwerx J. NAD(+) Metabolism and the Control of Energy Homeostasis: A Balancing Act between Mitochondria and the Nucleus. Cell Metab. 2015;22(1):31–53.

4. 4. Rajman L, Chwalek K, Sinclair DA. Therapeutic Potential of NAD-Boosting Molecules: The In Vivo Evidence. Cell Metab. 2018;27(3):529-47.

5. Covarrubias AJ, Perrone R, Grozio A, Verdin E. NAD(+) metabolism and its roles in cellular processes during ageing. Nat Rev Mol Cell Biol. 2021;22(2):119–41.

6. Katsyuba E, Romani M, Hofer D, Auwerx J. NAD(+) homeostasis in health and disease. Nat Metab. 2020;2(1):9–31.

7. Lopez-Otin C, Blasco MA, Partridge L, Serrano M, Kroemer G. The hallmarks of aging. Cell. 2013;153(6):1194–217.

8. Lopez-Otin C, Blasco MA, Partridge L, Serrano M, Kroemer G. Hallmarks of aging: An expanding universe. Cell. 2023;186(2):243–78.

9. Dutta S. CD38 as a driver of ageing. Nat Rev Endocrinol. 2026;22(3):137.

10. Camacho-Pereira J, Tarrago MG, Chini CCS, Nin V, Escande C, Warner GM, et al. CD38 Dictates Age-Related NAD Decline and Mitochondrial Dysfunction through an SIRT3-Dependent Mechanism. Cell Metab. 2016;23(6):1127–39.

11. Chini CCS, Peclat TR, Warner GM, Kashyap S, Espindola-Netto JM, de Oliveira GC, et al. CD38 ecto-enzyme in immune cells is induced during aging and regulates NAD(+) and NMN levels. Nat Metab. 2020;2(11):1284–304.

12. Imai SI. NAD World 3.0: the importance of the NMN transporter and eNAMPT in mammalian aging and longevity control. NPJ Aging. 2025;11(1):4.

13. Hope HC, de Sostoa J, Ginefra P, Andreatta M, Chiang YH, Ronet C, et al. Age-associated nicotinamide adenine dinucleotide decline drives CAR-T cell failure. Nat Cancer. 2025;6(9):1524–36.

14. Santiago-Cruz W, Espinosa E, Perez-Lara JC, Romero-Ramirez H, Devarajan P, Garcia-Garcia F, et al. CD38 promotes LPS-induced innate-like activation and proliferation of CD8(+) T lymphocytes in aged mice. Front Aging. 2025;6:1701685.

15. Clement J, Wong M, Poljak A, Sachdev P, Braidy N. The Plasma NAD(+) Metabolome Is Dysregulated in "Normal" Aging. Rejuvenation Res. 2019;22(2):121–30.

16. Massudi H, Grant R, Braidy N, Guest J, Farnsworth B, Guillemin GJ. Age-associated changes in oxidative stress and NAD+ metabolism in human tissue. PLoS One. 2012;7(7):e42357.

17. Chaleckis R, Murakami I, Takada J, Kondoh H, Yanagida M. Individual variability in human blood metabolites identifies age-related differences. Proc Natl Acad Sci U S A. 2016;113(16):4252–9.

18. Bagga P, Hariharan H, Wilson NE, Beer JC, Shinohara RT, Elliott MA, et al. Single-Voxel (1) H MR spectroscopy of cerebral nicotinamide adenine dinucleotide (NAD(+) ) in humans at 7T using a 32-channel volume coil. Magn Reson Med. 2020;83(3):806–14.

19. Cuenoud B, Ipek O, Shevlyakova M, Beaumont M, Cunnane SC, Gruetter R, et al. Brain NAD Is Associated With ATP Energy Production and Membrane Phospholipid Turnover in Humans. Front Aging Neurosci. 2020;12:609517.

20. Zhu XH, Lu M, Lee BY, Ugurbil K, Chen W. In vivo NAD assay reveals the intracellular NAD contents and redox state in healthy human brain and their age dependences. Proc Natl Acad Sci U S A. 2015;112(9):2876–81.

21. Migliavacca E, Tay SKH, Patel HP, Sonntag T, Civiletto G, McFarlane C, et al. Mitochondrial oxidative capacity and NAD(+) biosynthesis are reduced in human sarcopenia across ethnicities. Nat Commun. 2019;10(1):5808.

22. Kim LJ, Chalmers TJ, Madawala R, Smith GC, Li C, Das A, et al. Host-microbiome interactions in nicotinamide mononucleotide (NMN) deamidation. FEBS Lett. 2023;597(17):2196–220.

23. Shats I, Williams JG, Liu J, Makarov MV, Wu X, Lih FB, et al. Bacteria Boost Mammalian Host NAD Metabolism by Engaging the Deamidated Biosynthesis Pathway. Cell Metab. 2020;31(3):564–79 e7.

24. Ulanovskaya OA, Zuhl AM, Cravatt BF. NNMT promotes epigenetic remodeling in cancer by creating a metabolic methylation sink. Nat Chem Biol. 2013;9(5):300–6.

25. Conze DB, Crespo-Barreto J, Kruger CL. Safety assessment of nicotinamide riboside, a form of vitamin B(3). Hum Exp Toxicol. 2016;35(11):1149–60.

26. Yi L, Maier AB, Tao R, Lin Z, Vaidya A, Pendse S, et al. The efficacy and safety of beta-nicotinamide mononucleotide (NMN) supplementation in healthy middle-aged adults: a randomized, multicenter, double-blind, placebo-controlled, parallel-group, dose-dependent clinical trial. Geroscience. 2023;45(1):29–43.

27. Dollerup OL, Christensen B, Svart M, Schmidt MS, Sulek K, Ringgaard S, et al. A randomized placebo-controlled clinical trial of nicotinamide riboside in obese men: safety, insulin-sensitivity, and lipid-mobilizing effects. Am J Clin Nutr. 2018;108(2):343–53.

28. Dollerup OL, Chubanava S, Agerholm M, Sondergard SD, Altintas A, Moller AB, et al. Nicotinamide riboside does not alter mitochondrial respiration, content or morphology in skeletal muscle from obese and insulin-resistant men. J Physiol. 2020;598(4):731–54.

29. Dollerup OL, Trammell SAJ, Hartmann B, Holst JJ, Christensen B, Moller N, et al. Effects of Nicotinamide Riboside on Endocrine Pancreatic Function and Incretin Hormones in Nondiabetic Men With Obesity. J Clin Endocrinol Metab. 2019;104(11):5703–14.

30. Elhassan YS, Kluckova K, Fletcher RS, Schmidt MS, Garten A, Doig CL, et al. Nicotinamide Riboside Augments the Aged Human Skeletal Muscle NAD(+) Metabolome and Induces Transcriptomic and Anti-inflammatory Signatures. Cell Rep. 2019;28(7):1717–28 e6.

31. Pavlech LL, Yoon S, Gianturco SL, Storm KD, Yuen MV, Mattingly AN. Nicotinamide adenine dinucleotide: Summary Report. University of Maryland Center of Excellence in Regulatory Science and Innovation; 2021.

32. Kulikova V, Shabalin K, Nerinovski K, Dolle C, Niere M, Yakimov A, et al. Generation, Release, and Uptake of the NAD Precursor Nicotinic Acid Riboside by Human Cells. J Biol Chem. 2015;290(45):27124–37.

33. Nikiforov A, Kulikova V, Ziegler M. The human NAD metabolome: Functions, metabolism and compartmentalization. Crit Rev Biochem Mol Biol. 2015;50(4):284–97.

34. Hawkins J, Idoine R, Kwon J, Shao A, Dunne E, Hawkins E, et al. Randomized, placebo-controlled, pilot clinical study evaluating acute Niagen®+ IV and NAD+ IV in healthy adults. medRxiv. 2024:2024.06. 06.24308565.

35. Saqr AA, Kamali C, Brunnbauer P, Haep N, Koch P, Hillebrandt KH, et al. Optimized protocol for quantification of extracellular nicotinamide adenine dinucleotide: evaluating clinical parameters and pre-analytical factors for translational research. Front Med (Lausanne). 2023;10:1278641.

36. Vinten KT, Tretowicz MM, Coskun E, van Weeghel M, Canto C, Zapata-Perez R, et al. NAD(+) precursor supplementation in human ageing: clinical evidence and challenges. Nat Metab. 2025;7(10):1974–90.

37. Knop K, Hoogenboom R, Fischer D, Schubert US. Poly(ethylene glycol) in drug delivery: pros and cons as well as potential alternatives. Angew Chem Int Ed Engl. 2010;49(36):6288–308.

38. Zimmerman M, Kiefer R, Kerr S, Balling C. Reliability and validity of a self-report scale for daily assessments of the severity of anxiety symptoms. Compr Psychiatry. 2019;90:37–42.

39. Lovibond PF, Lovibond SH. The structure of negative emotional states: comparison of the Depression Anxiety Stress Scales (DASS) with the Beck Depression and Anxiety Inventories. Behav Res Ther. 1995;33(3):335–43.

40. Fisk JD, Doble SE. Construction and validation of a fatigue impact scale for daily administration (D-FIS). Qual Life Res. 2002;11(3):263–72.

41. Lawton KA, Berger A, Mitchell M, Milgram KE, Evans AM, Guo L, et al. Analysis of the adult human plasma metabolome. Pharmacogenomics. 2008;9(4):383–97.

42. Faul F, Erdfelder E, Lang AG, Buchner A. G*Power 3: a flexible statistical power analysis program for the social, behavioral, and biomedical sciences. Behav Res Methods. 2007;39(2):175–91.

43. Cohen J. Statistical power analysis for the behavioral sciences. 2nd ed. Hillsdale, N.J.: L. Erlbaum Associates; 1988. xxi, 567 p. p.

44. Galwey NW. A new measure of the effective number of tests, a practical tool for comparing families of non-independent significance tests. Genet Epidemiol. 2009;33(7):559–68.

45. Brydges CR. Effect Size Guidelines, Sample Size Calculations, and Statistical Power in Gerontology. Innov Aging. 2019;3(4):igz036.

46. Chen M, Yuan L, Chen B, Chang H, Luo J, Zhang H, et al. SLC29A1 and SLC29A2 are human nicotinamide cell membrane transporters. Nat Commun. 2025;16(1):1181.

47. Gasparrini M, Sorci L, Raffaelli N. Enzymology of extracellular NAD metabolism. Cell Mol Life Sci. 2021;78(7):3317–31.

48. Bruzzone S, Guida L, Zocchi E, Franco L, De Flora A. Connexin 43 hemi channels mediate Ca2+-regulated transmembrane NAD+ fluxes in intact cells. FASEB J. 2001;15(1):10–2.

49. Christen S, Redeuil K, Goulet L, Giner MP, Breton I, Rota R, et al. The differential impact of three different NAD(+) boosters on circulatory NAD and microbial metabolism in humans. Nat Metab. 2026;8(1):62–73.

50. McReynolds MR, Chellappa K, Chiles E, Jankowski C, Shen Y, Chen L, et al. NAD(+) flux is maintained in aged mice despite lower tissue concentrations. Cell Syst. 2021;12(12):1160–72 e4.

51. Conze D, Brenner C, Kruger CL. Safety and Metabolism of Long-term Administration of NIAGEN (Nicotinamide Riboside Chloride) in a Randomized, Double-Blind, Placebo-controlled Clinical Trial of Healthy Overweight Adults. Sci Rep. 2019;9(1):9772.

52. Morifuji M, Higashi S, Ebihara S, Nagata M. Ingestion of beta-nicotinamide mononucleotide increased blood NAD levels, maintained walking speed, and improved sleep quality in older adults in a double-blind randomized, placebo-controlled study. Geroscience. 2024;46(5):4671–88.

53. Yoshino M, Yoshino J, Kayser BD, Patti GJ, Franczyk MP, Mills KF, et al. Nicotinamide mononucleotide increases muscle insulin sensitivity in prediabetic women. Science. 2021;372(6547):1224–9.

54. Ye B, Pei Y, Wang L, Meng D, Zhang Y, Zou S, et al. NAD(+) supplementation prevents STING-induced senescence in CD8(+) T cells by improving mitochondrial homeostasis. J Cell Biochem. 2024;125(3):e30522.

55. Rutkowski B, Slominska E, Szolkiewicz M, Smolenski RT, Striley C, Rutkowski P, et al. N-methyl-2-pyridone-5-carboxamide: a novel uremic toxin? Kidney Int Suppl. 2003(84):S19–21.

56. Hikosaka K, Ikutani M, Shito M, Kazuma K, Gulshan M, Nagai Y, et al. Deficiency of nicotinamide mononucleotide adenylyltransferase 3 (nmnat3) causes hemolytic anemia by altering the glycolytic flow in mature erythrocytes. J Biol Chem. 2014;289(21):14796–811.

57. Pospieszna B, Kusy K, Slominska EM, Zielinski J, Ciekot-Soltysiak M. Erythrocyte nicotinamide adenine dinucleotide concentration is enhanced by systematic sports participation. BMC Sports Sci Med Rehabil. 2024;16(1):216.

