## Supplement for "Oral LNAD+ Rapidly Elevates Intracellular NAD and Metabolic Flux Without Elevating Circulating NAD: Evidence from a Randomized Controlled Trial"

Electronic Supplementary Materials

*Manuscript submitted for publication*

Sergey A. Kornilov, Waylon J. Hastings, Lynne Fahey McGrath, Michael Leitz-Langan, Andrew T. Magis, Steve M. Coppess, & Wendy Komac

### Contents

1 Supplementary Methods: Data Processing and Quality Control	4
  1.1 NAD+ Concentrations (icNAD, cirNAD)	4
  1.2 Clinical Laboratory Parameters	4
  1.3 Oxidative Stress and Inflammatory Biomarkers (ROM, hs-CRP, Beta-Galactosidase)	4
  1.4 NAD+ Metabolites (Metabolomics)	5
  1.5 Vital Signs	5
  1.6 Subjective Well-being	5
  1.7 Review of Systems (RoS)	6
  1.8 Wearable Activity and Sleep Monitoring (Fitbit)	6
  1.9 Multiplicity Correction	6

1.10 NAD+ Correlation Analysis…………………………………………………………………………………………………………………………………..6

List of Tables

Supplementary Table 1 Analysis populations.	3

Supplementary Table 2 Documented data corrections applied during quality control.	7

Supplementary Table 3 G;lobal Multiplicity correction: endpoint universe and unified M_eff.	7

Supplementary Table 4 ICNAD MMRM coefficient estimates (raw model parameterization).	9

Supplementary Table 5 CIRNAD MMRM coefficient estimates (raw model parameterization).	11

Supplementary Table 6 NAD+ Metabolites.	13

Supplementary Table 7 Baseline and Day 6 values for vitals and clinical laboratory tests.	14

Supplementary Table 8 Wearable (Fitbit) domain summary.	16

Supplementary Table 9 Review of Systems symptom incidence by body system.	17

Supplementary Table 10 Review of Systems body-system GLMM results.	19

Supplementary Table 11 Baseline Spearman correlations between NAD+ concentrations and clinical endpoints.	20

**List of Figures**

Supplementary Figure 1 CONSORT flow diagram.	3

Supplementary Figure 2 Sensitivity analysis (primary vs. full ITT cohort).	8

Supplementary Figure 3 Bootstrap distribution of the ICNAD treatment effect estimate (log2 scale).	9

Supplementary Figure 4 Bootstrap distribution of the ICNAD treatment effect estimate (raw scale).	10

Supplementary Figure 5 Bootstrap distribution of the CIRNAD treatment effect estimate (log2 scale).	11

Supplementary Figure 6 Bootstrap distribution of the CIRNAD treatment effect estimate (raw scale).	12

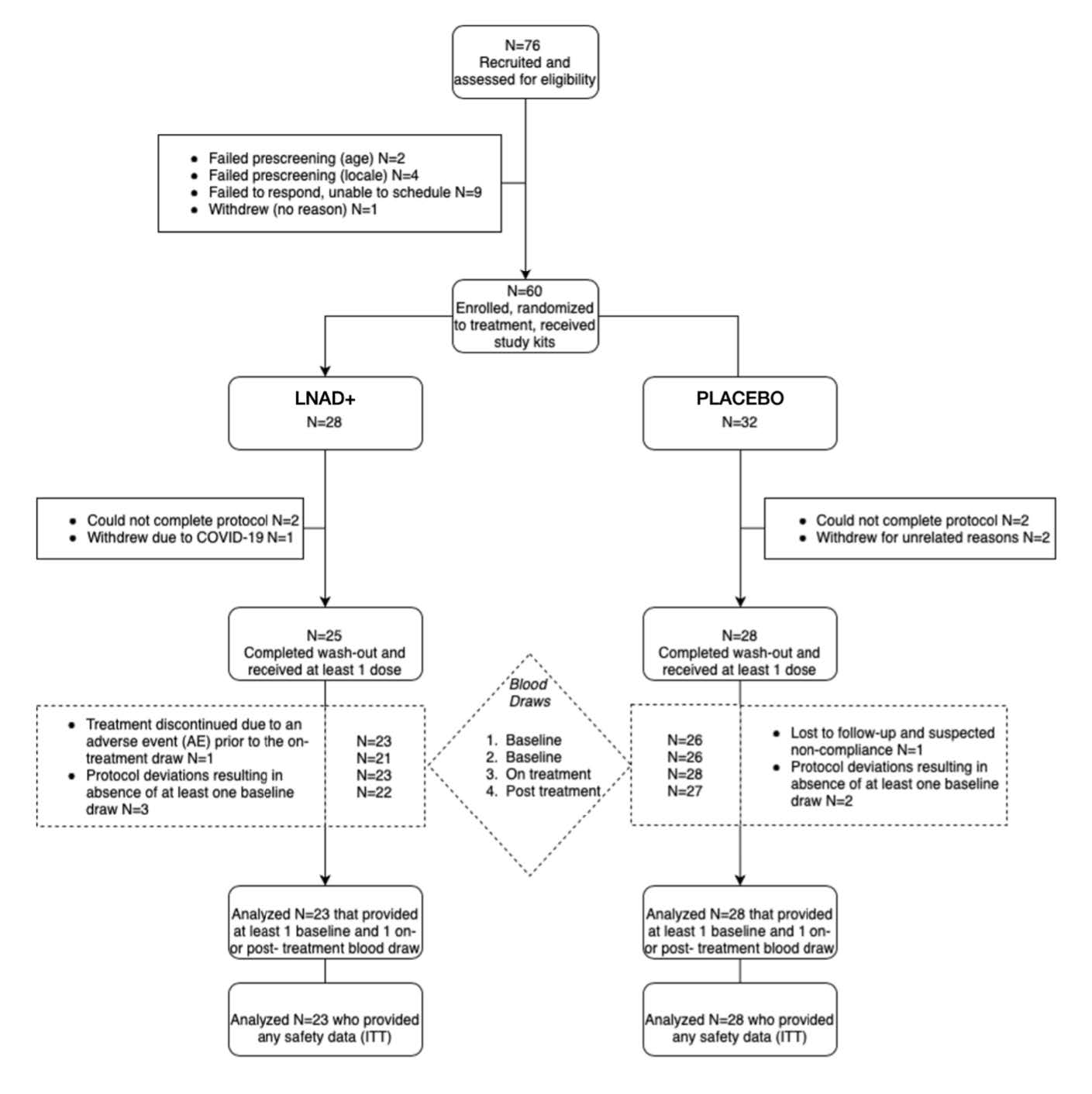

**Supplementary Figure 1.** CONSORT flow diagram. *Of 76 individuals recruited and assessed for eligibility, 60 were enrolled and randomized (LNAD+ N=28; Placebo N=32). Nine participants did not complete the protocol (could not complete: 4; withdrew: 3; COVID-19: 1; lost to follow-up with suspected non-compliance: 1). The final analysis included N=51 (ITT) and N=50 (primary analysis cohort, one placebo participant excluded post-hoc for icNAD decline consistent with recent B-vitamin supplement discontinuation).*

**Supplementary Table 1.** Analysis populations.

| **Population** | **N** | **Definition** |
| --- | --- | --- |
| Recruited | 76 | Assessed for eligibility |
| Randomized | 60 | Enrolled, randomized, received study kits |
| Completed | 51 | Completed washout, received ≥1 dose, provided ≥1 on/post-treatment draw |
| Full ITT | 51 | All completed participants (sensitivity analysis) |
| Primary | 50 | ITT minus one placebo participant excluded post-hoc for icNAD decline consistent with B-vitamin washout noncompliance |

*Mixed models used all available observed data under the missing-at-random assumption.*

### 1 Supplementary Methods: Data Processing and Quality Control

All analyses were implemented in R (v4.4.2) using a reproducible {targets} pipeline with computational environment locked via renv. The primary analytical model was MMRM implemented via the mmrm R package (v0.3.11) with REML estimation. Mixed models used all available observed data under the missing-at-random (MAR) assumption. Raw data provenance and processing steps are documented below for each data type.

**Visit schedule.** Two pre-treatment baseline blood draws were collected during the washout period (Baseline 1: days −4 to −3; Baseline 2: days −2 to −1 before treatment onset). On-treatment draws were collected at Day 4 (after 3 days of dosing) and Day 6 (1 day after the final dose). Patient-reported outcomes (DASS-21, CUXOS-D, D-FIS) were administered at three timepoints corresponding to Baseline 1 (day −7), Day 4, and Day 6. For clinical and NAD endpoints, baseline was defined as the average of the two pre-treatment measurements. For metabolomics, the most recent pre-treatment draw was used.

#### 1.1 NAD+ Concentrations (icNAD, cirNAD)

Intracellular NAD+ (icNAD) and circulating NAD+ (cirNAD) were measured as total NAD (NAD+ + NADH) via enzymatic cycling assay (Jinfiniti Precision Medicine). Values were extracted in their original units (icNAD in µM, cirNAD in nM) and log2-transformed for MMRM analysis. Baseline was computed as the average of two pre-treatment draws. Change from baseline (CHG) was computed on the log2 scale. The MMRM model was specified as: CHG = BASE + AVISIT

× TRT01A + SEX + AGE + BMIBL + VITDFL + SEX:AGE + us(AVISIT | USUBJID),

with Kenward–Roger denominator degrees of freedom. Nonparametric BCa bootstrap confidence intervals were computed using subject-level case resampling (1,000,000 replicates) with jackknife acceleration correction. Test–retest reliability across the two baseline draws was r = 0.95 (p < 0.001) for icNAD and r = 0.84 (p < 0.001) for cirNAD, supporting the use of averaged baselines to reduce measurement noise.

#### 1.2 Clinical Laboratory Parameters

Clinical laboratory data were derived from the raw wide-format source file, containing 52 analytes across liver function, proteins, glucose metabolism, lipid metabolism, inflammation, kidney func-tion, electrolytes, complete blood count (RBC, WBC, platelets), and oxidative stress panels. Data were pivoted to long format with standardized PARAMCD values. Documented data corrections were applied for specific participants with recording errors identified during QC (see Methods). Box–Cox power transformation was applied per parameter using car::powerTransform() to estimate optimal λ, with per-parameter overrides for known problematic distributions (log2 for albumin, MCH, MCHC, calcium, sodium, chloride, eGFR; log1p for basophils, basophil absolute, immature granulocytes, immature granulocytes absolute). Raw (untransformed) values were preserved for descriptive statistics. Baseline was computed as the average of Visits 1 and 2 on the transformed scale. The same MMRM model specification was applied as for NAD endpoints. Global M_eff/ Šidák correction was applied across all endpoint families (see Multiplicity Correction).

#### 1.3 Oxidative Stress and Inflammatory Biomarkers

Reactive oxygen metabolites (ROM; oxidative stress), high-sensitivity C-reactive protein (hs-CRP;

systemic inflammation), and beta-galactosidase (B-gal; senescence marker) were extracted from the raw NAD measurements data and analyzed alongside clinical laboratory parameters using the same Box–Cox transformation and MMRM framework.

#### 1.4 NAD+ Metabolites (Metabolomics)

Plasma metabolomics was performed by Metabolon Inc. using the Metabolon Global Discovery Panel. Data were received as median-normalized relative abundances for 1,043 identified metabo-lites in ADAM format. Five metabolic fate markers of primary interest (1-methylnicotinamide [MeNAM], N^1^-methyl-2-pyridone-5-carboxamide [2PY], nicotinamide [NAM], quinolinate, and trigonelline) were pre-specified based on the NAD+ catabolic pathway. Relative abundances were log2-transformed prior to modeling; metabolites with >50% missing values were excluded. No additional imputation was performed beyond vendor-level processing. Each metabolite was analyzed individually using the same MMRM specification as clinical endpoints (CHG = BASE + AVISIT × TRT01A + SEX + AGE + BMIBL + VITDFL + SEX:AGE, unstructured covariance, Kenward–Roger df). To stabilize variance estimates across the 1,000-feature metabolome, residual variances were moderated using empirical Bayes shrinkage (limma::squeezeVar), borrowing strength across features without shrinking coefficient estimates (MMRM-EB). This approach preserves the longitudinal covariance structure and baseline adjustment of MMRM while improving power for low-abundance metabolites. For the full metabolome ( 1,000 features), M_eff/Šidák correction was applied using the feature count as M_eff. The five pre-specified NAD fate markers were included in the global M_eff multiplicity framework (see Multiplicity Correction). Back-transformation from log2 scale to percent change was applied for reporting.

#### 1.5 Vital Signs

Vital signs (systolic BP, diastolic BP, heart rate) were derived from the raw source data. Duplicate readings per visit were averaged (e.g., Systolic 1 and Systolic 2). Documented data corrections were applied for recording errors identified during QC, including: decimal misplacement in weight recordings (2 participants), extra digit in systolic reading (1 participant), non-numeric character removal (1 participant), implausible weight changes corrected to baseline or interpolated values (2 participants), height measurement error (1 participant), a 3-way column rotation of HR/SBP/DBP values at one visit (1 participant, identified by SBP < DBP), and implausible diastolic BP spike at one visit (1 participant, readings of 134/176 mmHg against a typical range of 62–80 mmHg, corrected to average of baseline and final visit). BMI was always recalculated from corrected height and weight, never trusted from raw source. All corrections are documented in the analysis pipeline source code. Vital signs were analyzed on natural scales (no transformation) using the same MMRM framework.

#### 1.6 Subjective Well-being

Patient-reported outcomes were assessed using three validated instruments administered at baseline (Visit 1), Day 4 (Visit 2), and Day 6 (Visit 3):

- **DASS-21** (Depression Anxiety Stress Scales–21, adapted for daily administration): 21 items scored 0–3, yielding three 7-item subscale sum scores (range 0–21 each). Subscale item assignments follow Lovibond & Lovibond (1995): Depression items 3, 5, 10, 13, 16, 17, 21; Anxiety items 2, 4, 7, 9, 15, 19, 20; Stress items 1, 6, 8, 11, 12, 14, 18.
- **CUXOS-D** (Clinically Useful Anxiety Outcome Scale, Daily Version): 20-item self-report measure of anxiety severity scored 0–4 per item (total range 0–80; higher scores indicate greater anxiety).
- **D-FIS** (Daily Fatigue Impact Scale): 8-item measure of fatigue impact scored 0–4 per item (total range 0–32; higher scores indicate greater fatigue impact on daily functioning).

All scores were computed as total sums. No item-level missingness was observed (all completed assessments had fully endorsed items); two participants had missing assessments at Day 6 due to visit-level dropout. Internal consistency (Cronbach’s α) at baseline (N = 52): DASS-21 Depression α = 0.871, DASS-21 Stress α = 0.893, CUXOS-D α = 0.868, D-FIS α = 0.946. DASS-21 Anxiety

showed lower internal consistency (α = 0.483), attributable to floor effects in this healthy cohort (65% scored 0; mean 0.5 out of 21). Scores were analyzed on their natural scales (no transformation) using the same MMRM framework.

#### 1.7 Review of Systems (RoS)

A 71-item Review of Systems questionnaire was administered daily throughout the study period (Days 1–13). Each item was scored as present (1) or absent (0). Overall symptom incidence was analyzed as the proportion of symptom-days during the treatment period (Days 8–13) using a binomial generalized linear mixed model (GLMM) implemented in glmmTMB, with treatment-by-visit interaction as the fixed effect of interest and random intercepts for subjects. Wilson score 95% confidence intervals were computed for incidence proportions.

#### 1.8 Wearable Activity and Sleep Monitoring (Fitbit)

Continuous daily activity, sleep, and heart rate data were collected via Fitbit wearable devices throughout the study period. Twenty-one variables were analyzed across three domains: activity (daily steps, activity calories, calories out, lightly/fairly/very active minutes, sedentary minutes), sleep (total minutes asleep, time in bed, sleep efficiency, deep/REM/light/wake minutes, restless and awake counts), and heart rate (resting heart rate, cardio/fat burn/peak/out-of-range zone minutes). Quality control flags were applied: valid activity days required ≥600 minutes wear time

and ≥100 steps; valid sleep days required ≥180 minutes asleep; valid heart rate days required resting

HR between 30–120 bpm. Within-participant winsorization at 1st/99th percentiles was applied to reduce influence of extreme values. Yeo–Johnson power transformation was applied per variable to improve normality. Robust baseline estimates were computed as Huber M-estimator means over the 7-day washout period (Days 1–7), requiring at least 3 valid baseline days. Treatment-period data (Days 8–13) were analyzed using MMRM with adaptive covariance selection (compound symmetry, AR(1), heterogeneous AR(1), or unstructured, selected by AIC). The model included baseline, visit, treatment, visit-by-treatment interaction, age, sex, and baseline BMI as covariates. Satterthwaite denominator degrees of freedom were used. M_eff/Šidák correction was applied globally across all domains. Hedges’ g effect sizes were computed from the MMRM t-statistics for cross-variable comparison.

#### 1.9 Multiplicity Correction

M_eff/Šidák correction was the sole multiplicity adjustment. The effective number of independent tests (M_eff) was estimated across all 70 endpoints in the primary analysis pipeline (52 clinical laboratory, 6 vital sign, 5 wellbeing, 2 NAD, and 5 pre-specified NAD metabolites) by constructing a single pooled RBLW-shrunk correlation matrix of baseline values across all endpoints and applying the Galwey (2009) eigenvalue decomposition (Table S 3). This unified approach captures cross-domain correlations (e.g., between vital signs and lab parameters) that per-domain estimation would miss, yielding M_eff = 56 (20.0% reduction; shrinkage intensity λ = 0.044). Statistical signif-icance was evaluated against the global Šidák-adjusted threshold α = 1 − (1 − 0.05)1/56 ≈ 0.00092. Only icNAD and NAD catabolites MeNAM and 2PY survived this threshold.

### 1.10 NAD+ Correlation Analysis

To characterize the cross-sectional and longitudinal relationships between NAD+ concentrations and clinical endpoints, we performed exploratory bivariate Spearman rank correlation analyses between baseline icNAD or cirNAD values and baseline values of all clinical laboratory parameters (52 analytes spanning liver function, proteins, lipids, kidney function, electrolytes, hematology, inflammation, and oxida-tive stress), vital signs (6 parameters), subjective wellbeing scales (5 instruments), and NAD metabolites from untargeted metabolomics (nicotinamide [NAM], 1-methylnicotinamide [MeNAM], N^1^-methyl-2-pyridone-5-carboxamide [2PY], quinolinate, and trigonelline). Baseline correlations used data from all participants in the primary analysis cohort (N = 50) regardless of treatment assignment, as baseline values precede intervention. Baseline icNAD and cirNAD values were the average of two pre-treatment measurements (Baseline 1 and Baseline 2). Baseline values for clinical endpoints were derived from the prepared analysis data (BASE variable). For NAD metabolites, baseline values corresponded to the pre-treatment visit (AVISITN = 0) in the metabolomics ADAM dataset.

All correlations used pairwise complete observations, requiring a minimum of 5 complete pairs per test. P-values were computed using the asymptotic approximation for Spearman’s test. M_eff/Šidák correction was applied using the feature count as M_eff (135 tests). Only corrected p-values below 0.05 are considered statistically significant; nominal p-values below 0.05 without survival are reported as exploratory.

**Supplementary Table 2.** Documented data corrections applied during quality control.

| **Dataset** | **Visit** | **Variable** | **Raw Value** | **Corrected** | **Rationale** |
| --- | --- | --- | --- | --- | --- |
| Vitals | 2 | Weight | 1603.4 | 160.34 | Decimal misplacement (×10) |
| Vitals | 2 | Weight | 11.4 | 114 | Decimal misplacement (÷10) |
| Vitals | 2 | Systolic BP | 1471 | 147.1 | Extra digit |
| Vitals | 4 | Height | 65 | 60.5 | Measurement error (confirmed by adjacent visits) |
| Vitals | 2 | Systolic BP | “165+” | 165 | Non-numeric character removal |
| Vitals | 4 | Weight | 169 | 187 | 18 lb drop in 2 days; reverted to plausible value |
| Vitals | 2 | Weight | 163.8 | avg(v1,v3) | 28 lb jump in 1 day; interpolated |
| Vitals | 3 | Diastolic BP | 134 / 176 | 67.5 / 67.5 | Massive spike; subject range 62–80 mmHg; corrected to avg(BL, v4) |

*All corrections are implemented programmatically in the analysis pipeline source code (load_data.R) and are fully reproducible. BMI was always recalculated from corrected height and weight, never trusted from raw source.*

**Supplementary Table 3.** Global multiplicity correction: endpoint universe and unified M_eff.

| **Domain** | **Endpoints** |
| --- | --- |
| Clinical laboratory (52 analytes) | 52 |
| Vital signs (SBP, DBP, HR, MAP, BMI, Weight) | 6 |
| Wellbeing (DASS-21 ×3, CUXOS-D, D-FIS) | 5 |
| NAD (icNAD, cirNAD) | 2 |
| NAD metabolites (MeNAM, 2PY, NAM, quinolinate, trigonelline) | 5 |
| **Total endpoints** | **70** |
| **Unified M_eff (RBLW + Galwey)** | **56** |
| **Reduction** | **20.0%** |

*A single pooled RBLW-shrunk correlation matrix was constructed across all 70 baseline endpoint values and decomposed using the Galwey (2009) eigenvalue method. This captures cross-domain correlations that per-domain estimation would miss.*

Global Šidák threshold: α = 1 − (1 − 0.05)1/56 ≈ 0.000913.

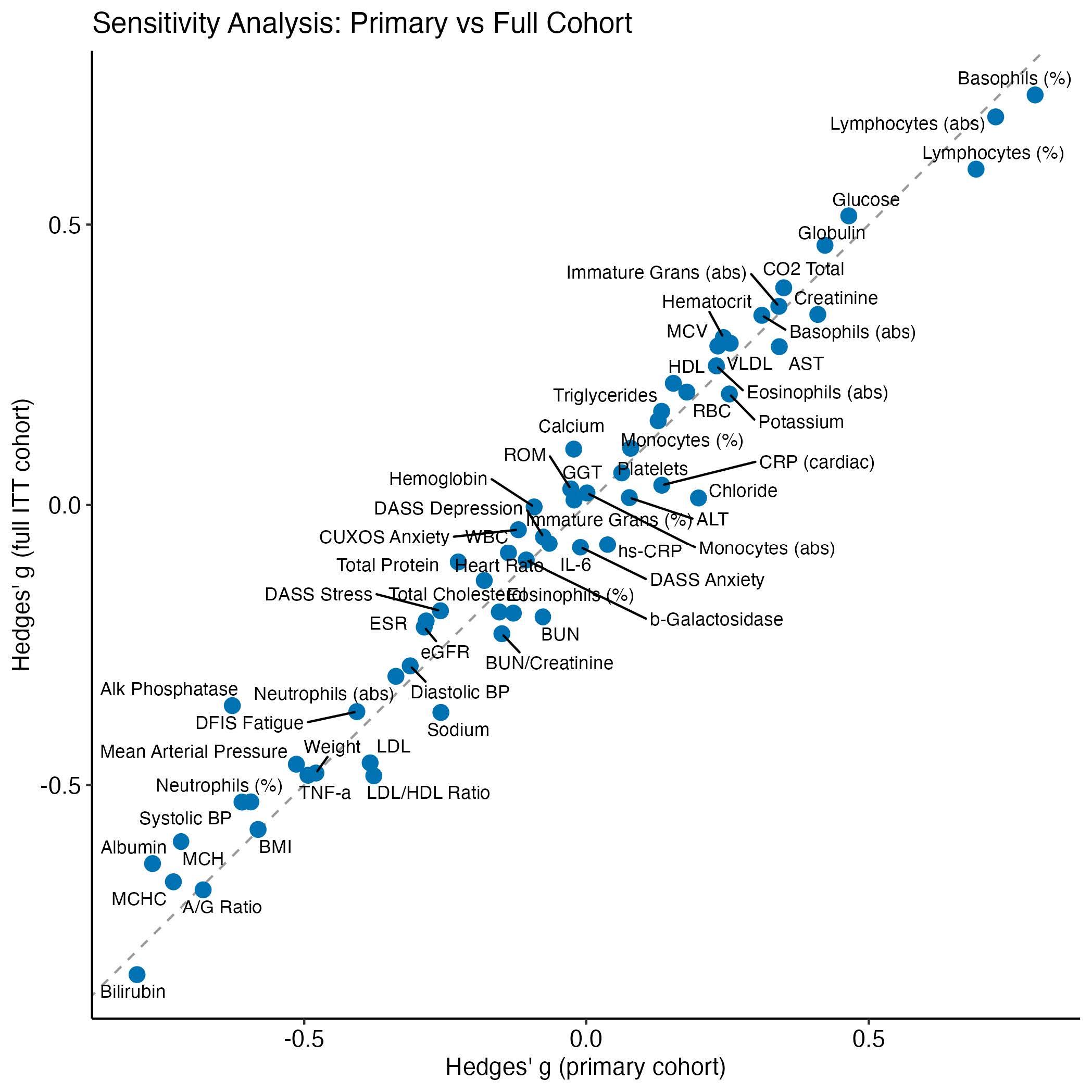

**Supplementary Figure 2** Sensitivity analysis (primary vs full ITT cohort). *Hedges' g effect size estimates for all endpoints in the primary analysis cohort (x-axis) versus the full intent-to-treat cohort (y-axis). Dashed line: identity. Proximity of points to the identity line demonstrates results are materially unchanged by the single exclusion*

**Supplementary Table 4.** ICNAD MMRM coefficient estimates (raw model parameterization).

| **Term** | **Estimate** | **Std. Error** | **df** | **t value** | **Pr(>\|t\|)** |
| --- | --- | --- | --- | --- | --- |
| (Intercept) | 1.8029 | 0.4936 | 42 | 3.6523 | 7.15e-04 |
| BASE | −0.2867 | 0.1011 | 42 | −2.8367 | 0.0070 |
| AVISITDay 6 | 0.1398 | 0.0216 | 46.0002 | 6.4632 | 5.84e-08 |
| TRT01APlacebo | −0.4669 | 0.0488 | 42.0004 | −9.5691 | 4.11e-12 |
| SEXM | 0.0068 | 0.0506 | 42.0001 | 0.1349 | 0.8933 |
| AGE | 0.0039 | 0.0046 | 42.0001 | 0.8538 | 0.3980 |
| BMIBL | −0.0035 | 0.0068 | 42.0001 | −0.5221 | 0.6044 |
| VITDFLY | 0.0518 | 0.0533 | 42.0002 | 0.9717 | 0.3368 |
| AVISITDay 6:TRT01APlacebo | −0.1461 | 0.0294 | 46.0002 | −4.9711 | 9.69e-06 |
| SEXM:AGE | −0.0033 | 0.0068 | 42.0001 | −0.4844 | 0.6306 |

*Reference levels: AVISIT = Day 4, TRT01A = LNAD. The TRT01APlacebo coefficient represents the treatment difference at Day 4; the Day 6 treatment contrast is TRT01APlacebo + AVISITDay6:TRT01APlacebo. Reported treatment effects in the* *manuscript use emmeans marginal contrasts at each visit. Kenward–Roger df. Primary analysis cohort (N = 50)*

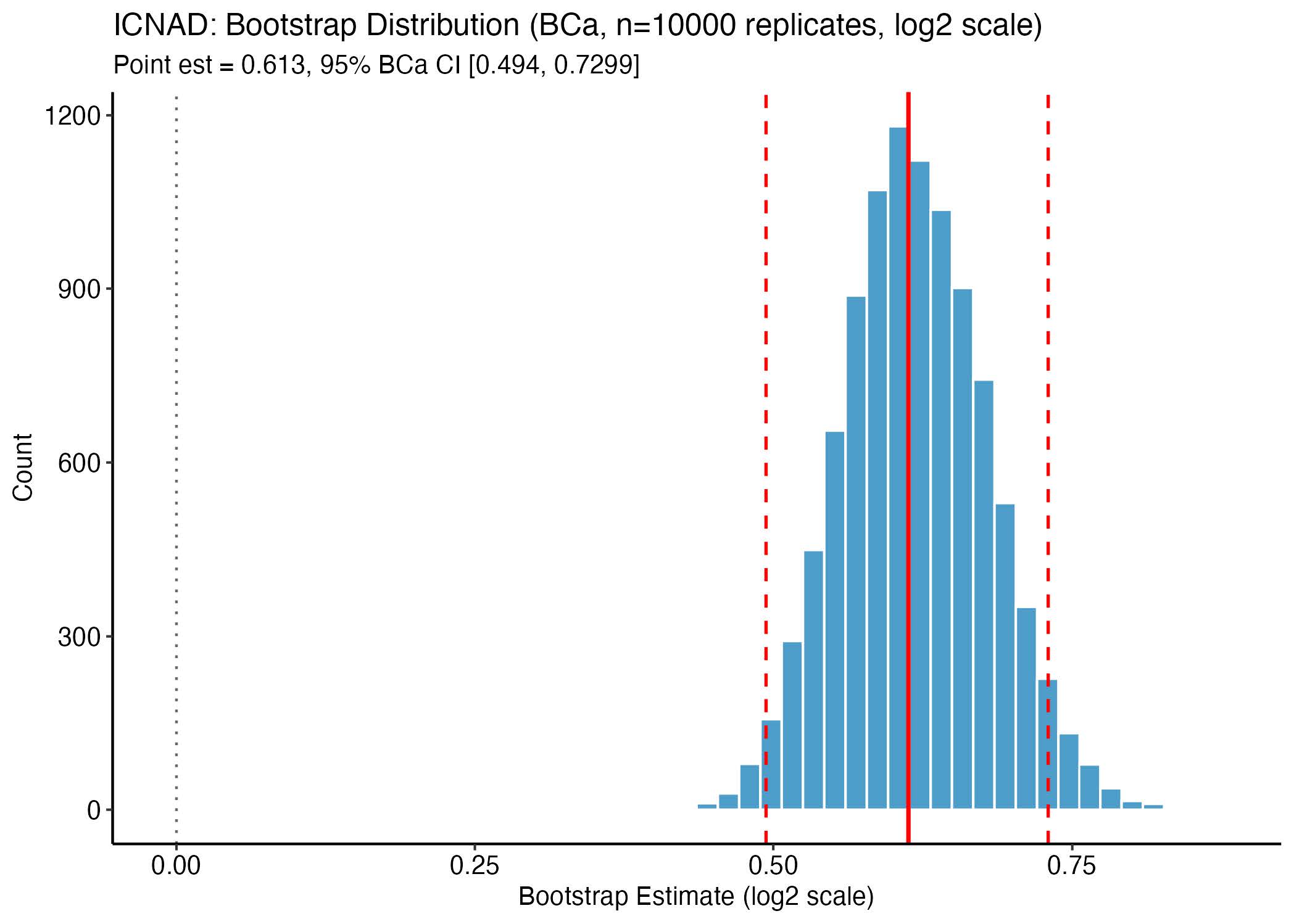

**Supplementary Figure 3.** Bootstrap distribution of the ICNAD treatment effect estimate (log2 scale). *Red solid line: point estimate. Red dashed lines: BCa 95% CI bounds. Gray dotted line: null (zero). BCa method with subject-**level case resampling*

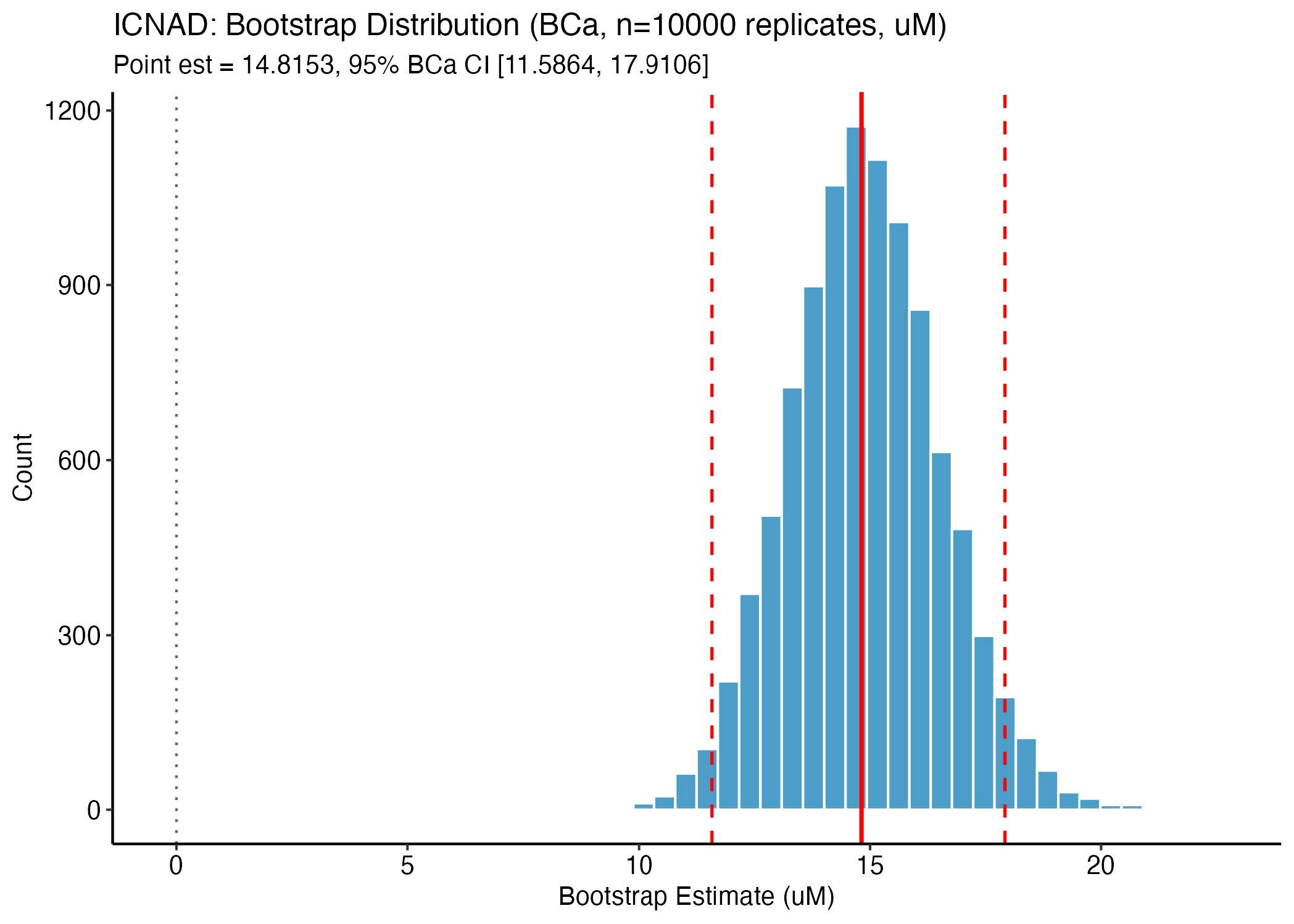

**Supplementary Figure 4.** Bootstrap distribution of the ICNAD treatment effect estimate (raw scale). *Red solid line: point estimate. Red dashed lines: BCa 95% CI bounds. Gray dotted line: null (zero). BCa method with subject-level case resampling*

**Supplementary Table 5.** CIRNAD MMRM coefficient estimates (raw model parameterization).

| **Term** | **Estimate** | **Std. Error** | **df** | **t value** | **Pr(>\|t\|)** |
| --- | --- | --- | --- | --- | --- |
| (Intercept) | 0.4008 | 0.5905 | 42.406 | 0.6788 | 0.5009 |
| BASE | −0.0448 | 0.0681 | 42.3449 | −0.6577 | 0.5143 |
| AVISITDay 6 | −0.0238 | 0.0516 | 46.0777 | −0.4611 | 0.6469 |
| TRT01APlacebo | −0.0538 | 0.0674 | 44.2895 | −0.7974 | 0.4294 |
| SEXM | 0.0138 | 0.0649 | 41.5607 | 0.213 | 0.8324 |
| AGE | −0.0012 | 0.0058 | 41.6944 | −0.2108 | 0.8341 |
| BMIBL | 0.0046 | 0.0087 | 41.6902 | 0.5306 | 0.5985 |
| VITDFLY | −0.0495 | 0.0685 | 42.0478 | −0.7227 | 0.4739 |
| AVISITDay 6:TRT01APlacebo | 0.0948 | 0.0701 | 46.0462 | 1.3516 | 0.1831 |
| SEXM:AGE | −0.0089 | 0.0085 | 41.8456 | −1.0381 | 0.3052 |

*Reference levels: AVISIT = Day 4, TRT01A = LNAD. The TRT01APlacebo coefficient represents the treatment difference at Day 4; the Day 6 treatment contrast is TRT01APlacebo + AVISITDay6:TRT01APlacebo. Reported treatment effects in the* *manuscript use emmeans marginal contrasts at each visit. Kenward–Roger df. Primary analysis cohort (N = 50)*

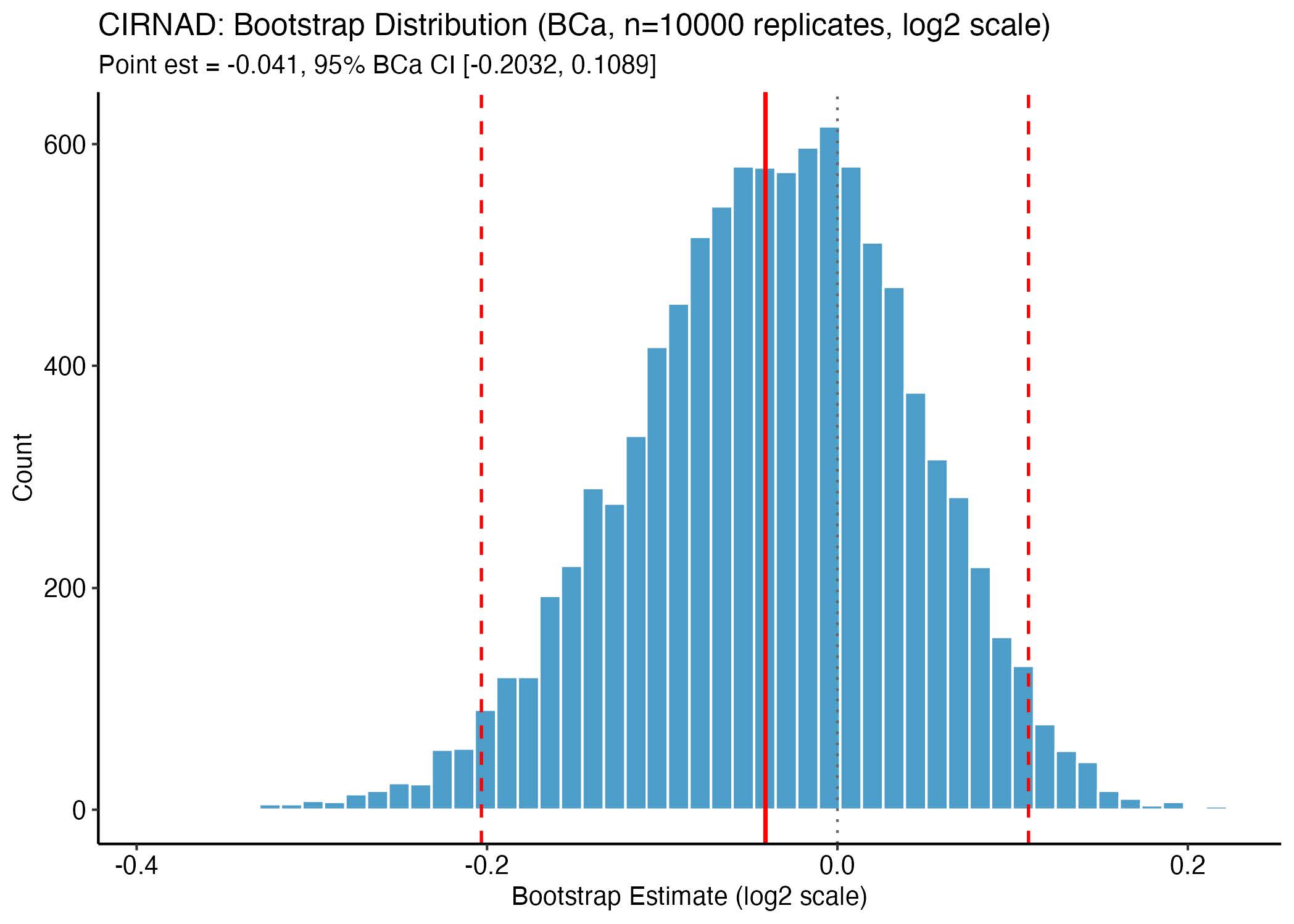

**Supplementary Figure 5.** Bootstrap distribution of the CIRNAD treatment effect estimate (log2 scale). *Red solid line: point estimate. Red dashed lines: BCa 95% CI bounds. Gray dotted line: null (zero). BCa method with subject-**level case resampling*

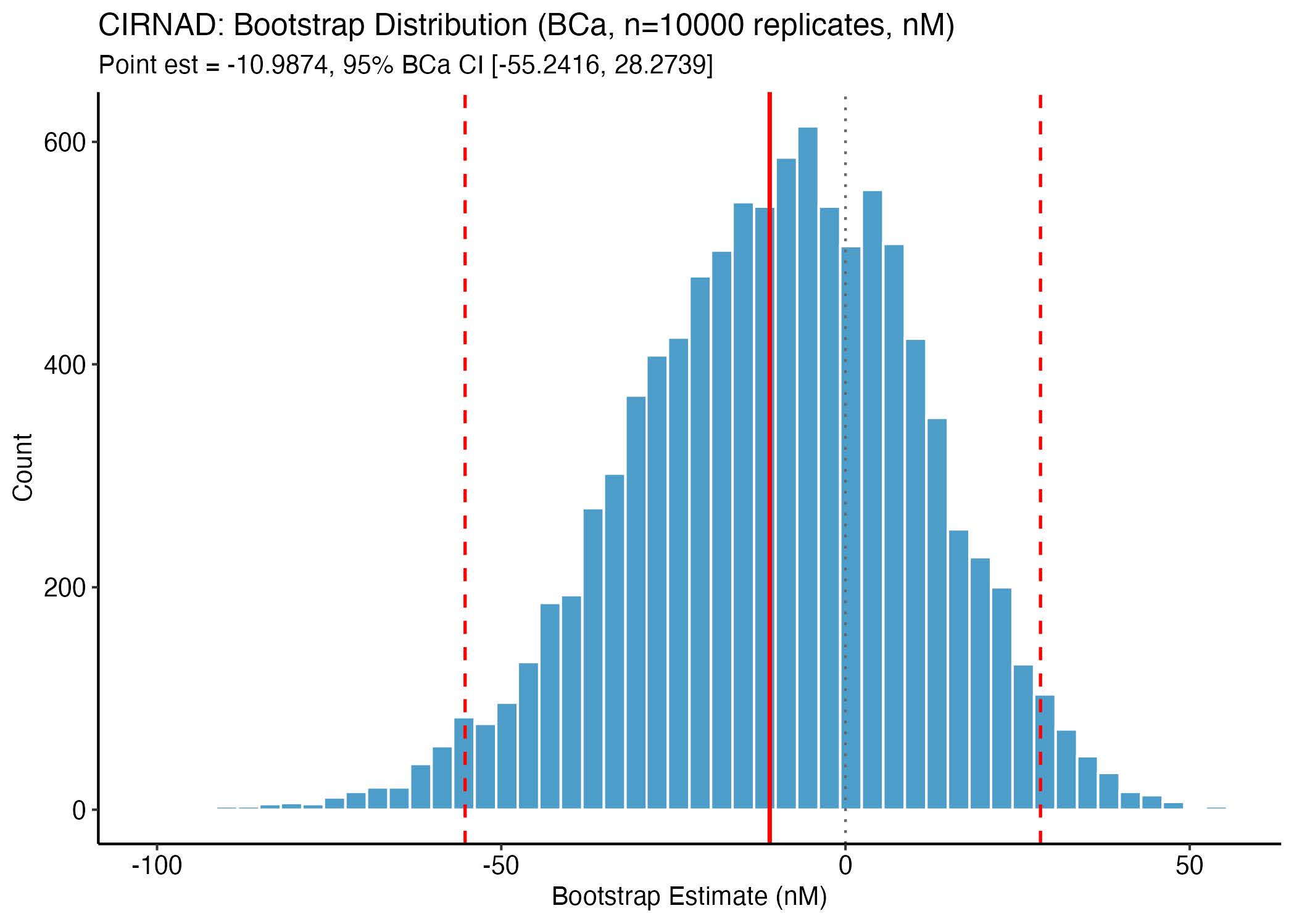

**Supplementary Figure 6.** Bootstrap distribution of the CIRNAD treatment effect estimate (raw scale). *Red solid line: point estimate. Red dashed lines: BCa 95% CI bounds. Gray dotted line: null (zero). BCa method with subject-level case resampling*

**Supplementary Table 6: NAD+ metabolites**

| **Analyte** | **Timepoint** | **Arm** | **N** | **Mean (SD)** | **SEM** | **95% CI** | **Change from BL** | **% Change (SD)** | **MMRM Est (%)** | **MMRM p** |
| --- | --- | --- | --- | --- | --- | --- | --- | --- | --- | --- |
| 1-Methylnicotinamide (MeNAM) | Baseline | LNAD | 23 | 0.72 (0.32) | 0.068 | [0.58, 0.86] |  | - | - | - |
| 1-Methylnicotinamide (MeNAM) | Baseline | Placebo | 26 | 0.73 (0.62) | 0.122 | [0.48, 0.98] |  | - | - | - |
| 1-Methylnicotinamide (MeNAM) | Day 4 | LNAD | 23 | 5.10 (2.59) | 0.540 | [3.98, 6.22] | 4.3775 | 724.0 (543.3) | 226.3 [171.9, 291.5] | 6.32e-17 |
| 1-Methylnicotinamide (MeNAM) | Day 4 | Placebo | 26 | 0.69 (0.31) | 0.061 | [0.57, 0.82] | -0.0384 | 26.9 (67.4) | 226.3 [171.9, 291.5] | 6.32e-17 |
| 1-Methylnicotinamide (MeNAM) | Day 6 | LNAD | 22 | 3.26 (1.88) | 0.400 | [2.43, 4.10] | 2.5457 | 401.7 (320.0) | 133.8 [94.8, 180.5] | 2.30e-12 |
| 1-Methylnicotinamide (MeNAM) | Day 6 | Placebo | 26 | 0.70 (0.54) | 0.106 | [0.48, 0.91] | -0.0359 | 8.5 (51.1) | 133.8 [94.8, 180.5] | 2.30e-12 |
| N1-Methyl-2-pyridone-5-carboxamide (2PY) | Baseline | LNAD | 23 | 0.69 (0.23) | 0.047 | [0.59, 0.78] |  | - | - | - |
| N1-Methyl-2-pyridone-5-carboxamide (2PY) | Baseline | Placebo | 26 | 0.66 (0.32) | 0.062 | [0.54, 0.79] |  | - | - | - |
| N1-Methyl-2-pyridone-5-carboxamide (2PY) | Day 4 | LNAD | 23 | 4.44 (1.53) | 0.318 | [3.78, 5.10] | 3.7566 | 631.6 (424.3) | 206.7 [166.1, 253.3] | 5.44e-20 |
| N1-Methyl-2-pyridone-5-carboxamide (2PY) | Day 4 | Placebo | 26 | 0.70 (0.29) | 0.057 | [0.58, 0.82] | 0.0330 | 19.7 (50.8) | 206.7 [166.1, 253.3] | 5.44e-20 |
| N1-Methyl-2-pyridone-5-carboxamide (2PY) | Day 6 | LNAD | 22 | 3.46 (1.19) | 0.254 | [2.93, 3.99] | 2.7710 | 478.4 (329.1) | 147.9 [115.1, 185.6] | 8.65e-17 |
| N1-Methyl-2-pyridone-5-carboxamide (2PY) | Day 6 | Placebo | 26 | 0.71 (0.33) | 0.065 | [0.57, 0.84] | 0.0429 | 19.0 (52.3) | 147.9 [115.1, 185.6] | 8.65e-17 |
| Nicotinamide (NAM) | Baseline | LNAD | 23 | 1.29 (0.57) | 0.118 | [1.04, 1.54] |  | - | - | - |
| Nicotinamide (NAM) | Baseline | Placebo | 26 | 1.02 (0.60) | 0.118 | [0.77, 1.26] |  | - | - | - |
| Nicotinamide (NAM) | Day 4 | LNAD | 23 | 2.19 (1.15) | 0.239 | [1.69, 2.68] | 0.8984 | 79.2 (91.3) | 28.5 [6.7, 54.6] | 0.009 |
| Nicotinamide (NAM) | Day 4 | Placebo | 26 | 1.22 (0.70) | 0.137 | [0.93, 1.50] | 0.1984 | 33.1 (72.6) | 28.5 [6.7, 54.6] | 0.009 |
| Nicotinamide (NAM) | Day 6 | LNAD | 22 | 1.68 (0.89) | 0.190 | [1.28, 2.07] | 0.3893 | 43.2 (101.1) | 11.9 [-7.1, 34.7] | 0.229 |
| Nicotinamide (NAM) | Day 6 | Placebo | 26 | 1.10 (0.57) | 0.113 | [0.87, 1.34] | 0.0864 | 29.1 (76.3) | 11.9 [-7.1, 34.7] | 0.229 |
| Quinolinate | Baseline | LNAD | 23 | 1.07 (0.43) | 0.090 | [0.89, 1.26] |  | - | - | - |
| Quinolinate | Baseline | Placebo | 26 | 1.11 (0.53) | 0.104 | [0.90, 1.33] |  | - | - | - |
| Quinolinate | Day 4 | LNAD | 23 | 1.05 (0.45) | 0.094 | [0.86, 1.25] | -0.0170 | 1.3 (28.4) | -4.5 [-14.4, 6.5] | 0.401 |
| Quinolinate | Day 4 | Placebo | 26 | 1.22 (0.59) | 0.116 | [0.99, 1.46] | 0.1119 | 22.1 (66.0) | -4.5 [-14.4, 6.5] | 0.401 |
| Quinolinate | Day 6 | LNAD | 22 | 1.05 (0.52) | 0.111 | [0.82, 1.28] | -0.0184 | -0.1 (35.8) | -0.9 [-11.1, 10.6] | 0.876 |
| Quinolinate | Day 6 | Placebo | 26 | 1.11 (0.49) | 0.096 | [0.91, 1.30] | -0.0056 | 15.5 (77.5) | -0.9 [-11.1, 10.6] | 0.876 |
| Trigonelline | Baseline | LNAD | 23 | 3.76 (3.33) | 0.694 | [2.33, 5.20] |  | - | - | - |
| Trigonelline | Baseline | Placebo | 26 | 3.06 (3.35) | 0.657 | [1.71, 4.42] |  | - | - | - |
| Trigonelline | Day 4 | LNAD | 23 | 4.11 (3.85) | 0.802 | [2.45, 5.77] | 0.3452 | 37.8 (101.8) | 13.6 [-14.4, 50.8] | 0.367 |
| Trigonelline | Day 4 | Placebo | 26 | 2.91 (2.79) | 0.547 | [1.78, 4.03] | -0.1576 | 31.3 (102.0) | 13.6 [-14.4, 50.8] | 0.367 |
| Trigonelline | Day 6 | LNAD | 22 | 4.05 (3.89) | 0.829 | [2.32, 5.77] | 0.2837 | 31.7 (114.0) | 15.0 [-13.3, 52.5] | 0.324 |
| Trigonelline | Day 6 | Placebo | 26 | 2.67 (2.76) | 0.542 | [1.55, 3.79] | -0.3949 | 38.1 (168.1) | 15.0 [-13.3, 52.5] | 0.324 |

*Metabolite values are Metabolon median-normalized relative abundance. MMRM Est (%) = treatment contrast (LNAD+ vs Placebo) on log2 scale, back-transformed to % change via (2d − 1) × 100. MeNAM = 1-methylnicotinamide; 2PY = N1-methyl-2-pyridone-5-carboxamide; NAM = nicotinamide. Primary analysis cohort.*

Supplementary Table 7: Baseline and Day 6 values for vitals and clinical laboratory tests

|  | LNAD+ Group | | | | Placebo Group | | | |  |  |  |
| --- | --- | --- | --- | --- | --- | --- | --- | --- | --- | --- | --- |
|  | Baseline | | Day 6 | | Baseline | | Day 6 | |  |  |  |
|  | Mean | SD | Mean | SD | Mean | SD | Mean | SD | Hedge’s g | p (nominal | p (M_eff) |
| A/G Ratio | 1.94 | 0.31 | 2.02 | 0.31 | 1.96 | 0.23 | 1.91 | 0.32 | -0.68 | 0.045 | 0.923 |
| Albumin | 4.46 | 0.28 | 4.52 | 0.28 | 4.52 | 0.22 | 4.45 | 0.32 | -0.77 | 0.033 | 0.843 |
| ALP | 75.15 | 20.09 | 75.20 | 19.58 | 72.72 | 21.91 | 71.18 | 19.98 | -0.63 | 0.067 | 0.98 |
| ALT | 22.04 | 14.45 | 20.16 | 8.61 | 22.02 | 11.11 | 21.27 | 11.51 | 0.08 | 0.796 | 1 |
| AST | 21.31 | 7.18 | 19.68 | 5.16 | 23.50 | 7.48 | 23.55 | 9.54 | 0.34 | 0.285 | 1 |
| B-gal | 287.24 | 176.29 | 310.42 | 210.79 | 354.52 | 309.73 | 295.09 | 298.52 |  |  |  |
| Baso (Abs.) | 0.04 | 0.05 | 0.04 | 0.05 | 0.05 | 0.04 | 0.05 | 0.05 | 0.31 | 0.329 | 1 |
| Baso (%) | 0.94 | 0.35 | 0.80 | 0.50 | 0.96 | 0.40 | 1.05 | 0.49 | 0.80 | 0.016 | 0.603 |
| TBILI | 0.41 | 0.27 | 0.46 | 0.39 | 0.44 | 0.17 | 0.36 | 0.18 | -0.80 | 0.019 | 0.654 |
| BUN | 15.76 | 4.68 | 15.60 | 4.40 | 16.17 | 4.79 | 15.95 | 4.57 | -0.08 | 0.83 | 1 |
| BUN/  Creatinine | 18.39 | 5.40 | 18.96 | 5.53 | 18.28 | 4.30 | 18.00 | 5.11 | -0.15 | 0.68 | 1 |
| CRP, Cardiac | 1.38 | 1.28 | 1.12 | 0.92 | 1.44 | 1.43 | 1.15 | 1.20 | 0.13 | 0.705 | 1 |
| Calcium | 9.43 | 0.31 | 9.47 | 0.43 | 9.35 | 0.36 | 9.40 | 0.34 | -0.02 | 0.95 | 1 |
| CO_2_ | 23.76 | 1.60 | 23.16 | 1.37 | 23.30 | 1.29 | 23.45 | 1.84 | 0.35 | 0.274 | 1 |
| Chloride | 101.83 | 1.62 | 102.04 | 1.88 | 102.26 | 1.41 | 102.50 | 1.57 | 0.20 | 0.528 | 1 |
| Cholesterol, Total | 209.39 | 47.65 | 210.24 | 46.03 | 212.04 | 44.95 | 209.91 | 41.92 | -0.15 | 0.652 | 1 |
| Creatinine | 0.86 | 0.14 | 0.83 | 0.11 | 0.89 | 0.19 | 0.90 | 0.17 | 0.41 | 0.138 | 1 |
| eGFR | 88.76 | 12.20 | 90.84 | 11.36 | 87.74 | 15.21 | 86.59 | 13.72 | -0.29 | 0.247 | 1 |
| Eos (Abs.) | 0.14 | 0.09 | 0.13 | 0.12 | 0.14 | 0.10 | 0.15 | 0.10 | 0.23 | 0.422 | 1 |
| Eos (%) | 2.54 | 1.49 | 2.52 | 1.94 | 2.67 | 1.66 | 2.73 | 2.03 | -0.13 | 0.612 | 1 |
| GGT | 19.48 | 16.31 | 16.12 | 12.21 | 23.46 | 18.96 | 21.73 | 16.68 | 0.06 | 0.860 | 1 |
| Globulin, Total | 2.35 | 0.29 | 2.28 | 0.30 | 2.35 | 0.27 | 2.37 | 0.32 | 0.42 | 0.193 | 1 |
| Glucose | 96.63 | 18.15 | 93.28 | 23.23 | 95.89 | 17.90 | 101.23 | 25.53 | 0.46 | 0.198 | 1 |
| HDL Cholesterol | 63.28 | 20.68 | 62.96 | 22.26 | 65.78 | 17.46 | 66.23 | 20.99 | 0.15 | 0.673 | 1 |
| Hematocrit | 41.07 | 3.49 | 40.18 | 3.11 | 41.68 | 2.71 | 41.15 | 2.88 | 0.24 | 0.446 | 1 |
| Hemoglobin | 13.86 | 1.28 | 13.69 | 1.16 | 14.06 | 0.93 | 13.88 | 0.94 | -0.09 | 0.758 | 1 |
| Grans (Abs) | 0.00 | 0.00 | 0.00 | 0.00 | 0.00 | 0.00 | 0.00 | 0.02 | 0.34 | 0.346 | 1 |
| Grans (%) | 0.06 | 0.21 | 0.08 | 0.28 | 0.04 | 0.14 | 0.09 | 0.29 | -0.02 | 0.946 | 1 |
| IL-6 | 2.64 | 0.65 | 2.63 | 0.92 | 3.90 | 3.09 | 3.41 | 3.11 | -0.07 | 0.866 | 1 |
| LDL | 124.17 | 35.68 | 124.62 | 31.66 | 125.15 | 40.30 | 122.14 | 34.05 | -0.38 | 0.288 | 1 |
| LDL/HDL Ratio | 2.16 | 0.94 | 2.09 | 0.77 | 2.05 | 0.91 | 1.98 | 0.76 | -0.38 | 0.288 | 1 |
| Lymphs (Abs) | 1.66 | 0.49 | 1.56 | 0.47 | 1.80 | 0.58 | 1.89 | 0.50 | 0.73 | 0.037 | 0.876 |
| Lymphs (%) | 30.44 | 5.88 | 29.32 | 7.30 | 33.02 | 6.13 | 35.00 | 7.26 | 0.69 | 0.053 | 0.953 |
| MCH | 30.15 | 1.43 | 30.44 | 1.48 | 30.43 | 1.31 | 30.44 | 1.28 | -0.72 | 0.031 | 0.826 |
| MCHC | 33.73 | 0.67 | 34.07 | 0.76 | 33.74 | 0.50 | 33.74 | 0.78 | -0.73 | 0.028 | 0.80 |
| MCV | 89.41 | 3.62 | 89.32 | 3.59 | 90.22 | 3.20 | 90.32 | 3.34 | 0.23 | 0.421 | 1 |
| Monocytes (Abs) | 0.46 | 0.12 | 0.45 | 0.13 | 0.48 | 0.18 | 0.48 | 0.16 | 0.00 | 0.997 | 1 |
| Monocytes (%) | 8.59 | 1.85 | 8.48 | 2.06 | 8.70 | 2.05 | 8.82 | 1.97 | 0.13 | 0.687 | 1 |
| Neutrophils (Abs) | 3.16 | 0.88 | 3.15 | 0.80 | 3.07 | 1.00 | 3.01 | 1.35 | -0.34 | 0.334 | 1 |
| Neutrophils | 57.43 | 7.12 | 58.80 | 8.76 | 54.61 | 7.17 | 52.32 | 8.93 | -0.59 | 0.093 | 0.996 |
| Platelets | 255.04 | 60.82 | 265.12 | 63.79 | 269.70 | 59.40 | 266.82 | 64.28 | 0.08 | 0.809 | 1 |
| Potassium | 4.34 | 0.29 | 4.35 | 0.29 | 4.31 | 0.28 | 4.36 | 0.38 | 0.25 | 0.477 | 1 |
| Protein, Total | 6.81 | 0.35 | 6.80 | 0.41 | 6.88 | 0.35 | 6.82 | 0.40 | -0.23 | 0.505 | 1 |
| RBC | 4.60 | 0.41 | 4.50 | 0.39 | 4.63 | 0.37 | 4.57 | 0.36 | 0.18 | 0.573 | 1 |
| Sedimentation Rate | 7.80 | 7.55 | 7.64 | 5.35 | 7.61 | 9.79 | 7.05 | 6.51 | -0.28 | 0.412 | 1 |
| Sodium | 139.39 | 1.09 | 139.52 | 1.83 | 139.65 | 1.16 | 139.50 | 1.79 | -0.26 | 0.417 | 1 |
| Triglycerides | 118.04 | 127.39 | 153.20 | 320.29 | 116.63 | 72.80 | 120.18 | 89.46 | 0.13 | 0.700 | 1 |
| TNF-a | 0.94 | 0.32 | 0.95 | 0.30 | 9.13 | 38.70 | 11.20 | 47.93 | -0.49 | 0.127 | 1 |
| VLDL Cholesterol | 21.94 | 25.82 | 16.17 | 6.84 | 21.11 | 12.91 | 21.55 | 15.44 | 0.25 | 0.489 | 1 |
| WBC | 5.45 | 1.30 | 5.34 | 1.09 | 5.56 | 1.55 | 5.59 | 1.77 | -0.14 | 0.687 | 1 |

*NOTE: *p- Hedges' g effect size and nominal MMRM p-value for the Day 6 treatment contrast (LNAD+ vs Placebo). p (M_eff) = global Šidák-corrected p-value (M_eff = 55). Primary analysis cohort (N = 50).*

*A/G= albumin to globulin ratio; ALP = Alkaline phosphatase; ALT = Alanine Transaminase; AST = Aspartate Transaminase; TBILI = total bilirubin; BUN = Blood Urea Nitrogen; CRP = C-reactive protein; DBP = Diastolic blood pressure; eGRF = Estimated Glomerular Filtration Rate; GGT = Gamma Glutamyl Transferase; HDL = High Density Lipoprotein; IL-6 = Interleukin 6; LDL = Low Density Lipoprotein; MCH = Mean Corpuscular Hemoglobin; MCHC = Mean Corpuscular Hemoglobin Concentration; MCV = Mean Corpuscular Volume; RBC = Red Blood Cell Count; SBP = Systolic Blood Pressure; SD = Standard Deviation; TNF-a = Tumor Necrosis Factor Alpha; WBC = White Blood Cell Count*

**Supplementary Table 8.** Wearable (Fitbit) domain summary.

| **domain** | **AVISIT** | **n_vars** | **n_sig** | **mean_effect** | **min_p** |
| --- | --- | --- | --- | --- | --- |
| activity | Day8 | 7 | 0 | 4.256139 | 0.05505 |
| activity | Day9 | 7 | 0 | 5.220002 | 0.148913 |
| activity | Day10 | 7 | 0 | 5.845532 | 0.065186 |
| activity | Day11 | 7 | 0 | 5.027732 | 0.061998 |
| activity | Day12 | 7 | 0 | 3.85194 | 0.054462 |
| activity | Day13 | 7 | 3 | 10.269809 | 0.030492 |
| heart_rate | Day8 | 5 | 2 | −271.531122 | 0.008247 |
| heart_rate | Day9 | 5 | 0 | −142.490044 | 0.201278 |
| heart_rate | Day10 | 5 | 0 | −79.143451 | 0.253014 |
| heart_rate | Day11 | 5 | 1 | −35.764855 | 0.021456 |
| heart_rate | Day12 | 5 | 0 | −122.448949 | 0.368089 |
| heart_rate | Day13 | 5 | 0 | 258.663098 | 0.083342 |
| sleep | Day8 | 9 | 0 | 17326041291093 | 0.055838 |
| sleep | Day9 | 9 | 3 | 32990760652388.8 | 0.000681 |
| sleep | Day10 | 9 | 0 | −6374379898449.68 | 0.065893 |
| sleep | Day11 | 9 | 0 | −4082799285103.81 | 0.250268 |
| sleep | Day12 | 9 | 0 | −2584557951274.76 | 0.113585 |
| sleep | Day13 | 9 | 0 | 34882430027398.2 | 0.251315 |

*Per-domain summary of variables analyzed, quality control filtering, and sample sizes after exclusions.*

**Supplementary Table 9.** Review of Systems symptom incidence by body system.

| **Body System** | **Symptom** | **LNAD+ (%)** | | **Placebo (%)** |
| --- | --- | --- | --- | --- |
| Allergy | itchy-eyes | 9.9 | | 2.8 |
|  | runny-nose | 17.4 | | 6.6 |
|  | sneezing | 11.2 | | 3.3 |
| Cardiovascular | chest-pain | 0 | | 0 |
|  | difficulty-breathing-sleep | 0 | | 0 |
|  | leg-pain | 0 | | 0 |
|  | leg-swelling | 0 | | 0 |
|  | palpitations | 1.2 | | 0 |
|  | sob-when-lying-down | 0 | | 0 |
| Constitution | chills | | 0.6 | 0 |
|  | fever | | 0.6 | 0 |
|  | malaise-fatigue | | 10.6 | 2.8 |
|  | sweating | | 1.2 | 1.7 |
|  | weakness-consitution | | 3.7 | 2.2 |
|  | weight-loss | | 0.6 | 0 |
| Endocrine-Hematology | easy-bruising-bleeding | | 0 | 0 |
|  | increased-thirst | | 1.9 | 1.7 |
| Eyes | blurred-vision | | 0 | 2.2 |
|  | double-vision | | 0 | 0 |
|  | eye-discharge | | 1.2 | 5 |
|  | eye-pain | | 1.9 | 0 |
|  | eye-redness | | 2.5 | 2.8 |
|  | light-sensitivity | | 0.6 | 1.7 |
| Gastrointestinal | abdominal-pain | | 1.2 | 0 |
|  | black-tarry-stool | | 0 | 0 |
|  | blood-in-stool | | 0 | 0 |
|  | constipation | | 6.2 | 1.7 |
|  | diarrhea | | 3.1 | 3.3 |
|  | heartburn | | 3.1 | 0 |
|  | nausea | | 1.2 | 1.7 |
|  | vomiting | | 0 | 0 |
| Genitourinary | blood-urine | | 0.6 | 0 |
|  | flank-pain | | 0 | 0 |
|  | urinary-frequency | | 5.6 | 5 |
|  | urinary-pain | | 0 | 2.2 |
|  | urinary-urgency | | 5 | 1.1 |
| Head-Ear-Nose-Throat | congestion | | 6.2 | 1.7 |
|  | ear-pain | | 0 | 0 |
|  | hearing-loss | | 0 | 0 |
|  | noisy-breathing | | 0 | 0.6 |
|  | nosebleeds | | 0 | 0 |
|  | ringing-ears | | 7.5 | 8.8 |
|  | sinus-pain | | 3.1 | 0.6 |
|  | sore-throat | | 3.7 | 2.8 |
| Musculoskeletal | back-pain | | 3.7 | 14.9 |
|  | falls | | 0 | 0 |
|  | joint-pain | | 11.2 | 15.5 |
|  | muscle-pain | | 6.2 | 8.3 |
|  | neck-pain | | 6.2 | 6.1 |
| Neurological | dizziness | | 1.2 | 0.6 |
|  | headaches | | 10.6 | 10.5 |
|  | loss-of-consciousness | | 0 | 0 |
|  | seizures | | 0 | 0 |
|  | sensory-change | | 0.6 | 0 |
|  | speech-change | | 0 | 0 |
|  | tingling | | 9.9 | 0 |
|  | tremor | | 0 | 0 |
|  | weakness-neurological | | 5.6 | 2.2 |
| Psychiatric | anxiety | | 3.7 | 15.5 |
|  | difficulty-sleeping | | 11.8 | 17.7 |
|  | memory-loss | | 0 | 0.6 |
|  | panic-attacks | | 0 | 0 |
|  | sadness-low-mood | | 6.2 | 12.2 |
|  | sleeping-too-much | | 3.1 | 0 |
| Respiratory | cough | | 5.6 | 2.8 |
|  | coughing-blood | | 0 | 0 |
|  | coughing-phlegm | | 5 | 1.7 |
|  | shortness-of-breath | | 1.9 | 0 |
|  | wheezing | | 1.9 | 0 |
| Skin | itching | | 1.2 | 0 |
|  | rash | | 0 | 0 |

*Values are percentage of participant-days on which the symptom was endorsed during the treatment period (Days 8–13). Overall treatment OR = 1.6 (GLMM p = 0.43). ROS data derived from NADStudy.Assess.FINAL.txt (71 binary symptoms, 7 daily* *assessments). Body system groupings from study metadata. Primary analysis cohort (N = 50)*

**Supplementary Table 10.** Review of Systems body-system GLMM results.

| **Body System** | **Odds Ratio** | **95% CI** | **p-value** | **Min Events** | **Convergence** |
| --- | --- | --- | --- | --- | --- |
| Allergy | 34.14 | (0.09, 144.86) | 0.506 | 14 | good |
| Constitution | 9.16 | (0.41, 11.90) | 0.354 | 10 | good |
| Endocrine-Hematology | 4.89 | (0.03, 88.07) | 0.822 | 3 | good |
| Eyes | 2.04 | (0.03, 15.73) | 0.831 | 8 | good |
| Gastrointestinal | 7.56 | (0.42, 9.64) | 0.376 | 12 | good |
| Genitourinary | 3.6 | (0.11, 15.52) | 0.846 | 11 | good |
| Head-Ear-Nose-Throat | 1.8 | (0.01, 23.19) | 0.777 | 21 | good |
| Musculoskeletal | 1.44 | (0.00, 27.32) | 0.649 | 31 | good |
| Neurological | 4.83 | (0.21, 11.74) | 0.657 | 22 | good |
| Psychiatric | 8.2 | (0.18, 24.77) | 0.554 | 34 | good |
| Respiratory | 10.24 | (0.00, 1103.32) | 0.788 | 6 | good |

*Binary GLMM (glmmTMB) odds ratios for symptom incidence (LNAD+ vs Placebo) during treatment period (Days 8–13), with random intercepts for subjects. Covariates: AGE, SEX. Min Events = minimum symptom-positive participant-days per arm.* *No body system reached nominal significance. Primary analysis cohort (N = 50)*

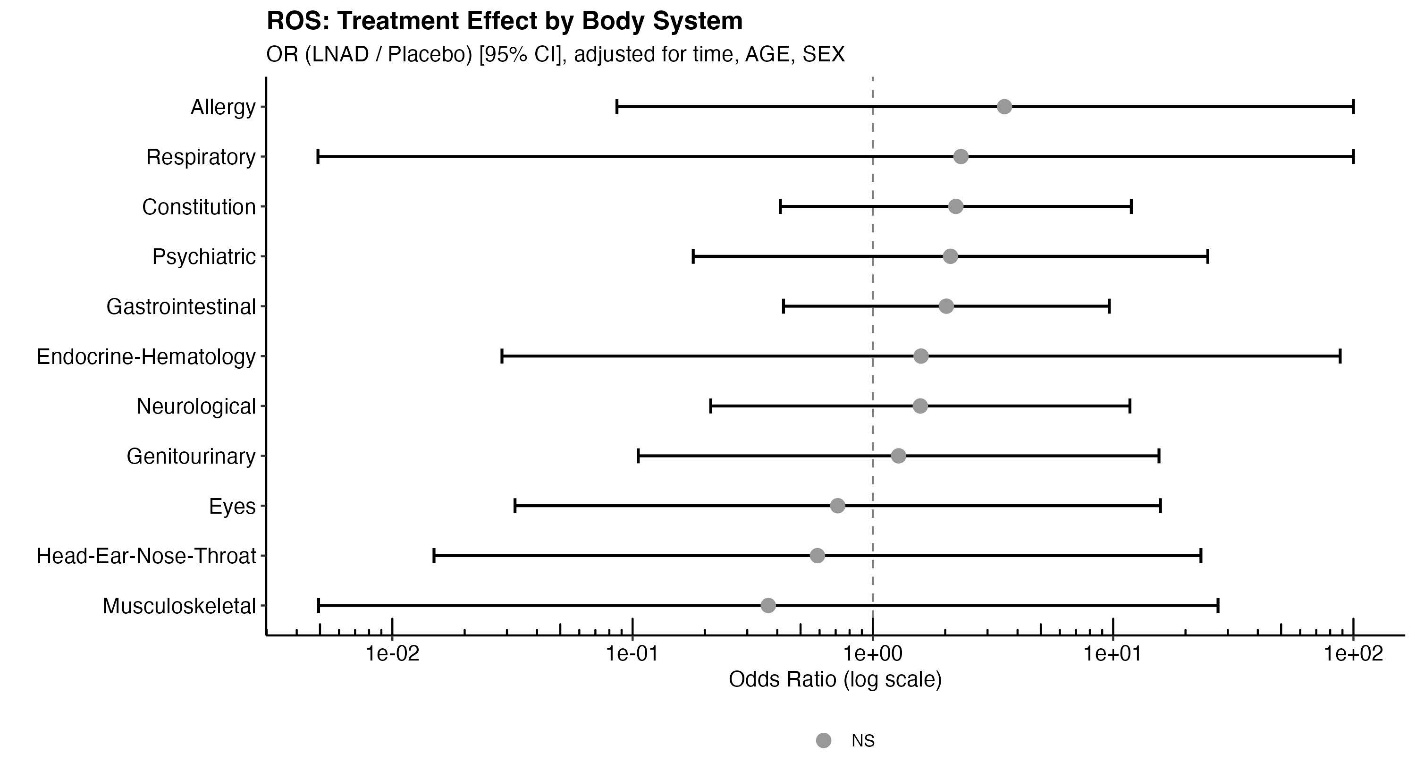

**Supplementary Figure 7.** Review of Systems body-system GLMM forest plot. *GLMM odds ratios (LNAD+ vs Placebo) for symptom incidence by body system during the treatment period. Squares: point estimates; horizontal bars: 95% confidence intervals. No body system showed a statistically significant difference. Primary analysis cohort*

**Supplementary Table 11.** Baseline Spearman correlations between NAD+ concentrations and clinical endpoints.

| **NAD Measure** | **Domain** | **Endpoint** | **rho** | **p (nominal)** | **p (adj)** | **n** |
| --- | --- | --- | --- | --- | --- | --- |
| **cirNAD** | **labs** | **TNF-a** | **−0.56** | **2.84e-05** | **0.003** | **50** |
|  | **labs** | **CREATININE** | **−0.51** | **1.43e-04** | **0.006** | **50** |
|  | **labs** | **HDL CHOLESTEROL** | **0.43** | **1.87e-03** | **0.042** | **50** |
|  | labs | EOS | −0.4 | 4.16e-03 | 0.08 | 50 |
|  | labs | ALKALINEPHOSPHATASE | −0.39 | 4.75e-03 | 0.08 | 50 |
|  | nad_metabolites | Quinolinate | −0.37 | 8.69e-03 | 0.106 | 49 |
|  | labs | SEDIMENTATIONRATEWEST-ERGREN | −0.37 | 8.88e-03 | 0.106 | 50 |
|  | vitals | WEIGHT | −0.36 | 1.14e-02 | 0.111 | 50 |
|  | labs | EOSABSOLUTE | −0.35 | 1.16e-02 | 0.111 | 50 |
|  | labs | VLDLCHOLESTEROLCAL | −0.34 | 1.68e-02 | 0.132 | 50 |
|  | labs | GLUCOSE | −0.33 | 1.94e-02 | 0.145 | 50 |
|  | labs | EGFR | 0.32 | 2.56e-02 | 0.18 | 50 |
|  | labs | BGAL | −0.31 | 2.79e-02 | 0.187 | 50 |
|  | nad_metabolites | MeNAM | 0.31 | 3.07e-02 | 0.196 | 49 |
|  | labs | INTERLEUKIN6SERUM | −0.3 | 3.24e-02 | 0.197 | 50 |
|  | labs | TRIGLYCERIDES | −0.3 | 3.61e-02 | 0.206 | 50 |
|  | vitals | DIABP | −0.28 | 4.83e-02 | 0.24 | 50 |
| **icNAD** | **labs** | **HEMOGLOBIN** | **0.54** | **5.03e-05** | **0.003** | **50** |
|  | **labs** | **HEMATOCRIT** | **0.5** | **1.87e-04** | **0.006** | **50** |
|  | **nad_metabolites** | **NAM** | **0.47** | **6.64e-04** | **0.018** | **49** |
|  | labs | MCH | 0.37 | 8.86e-03 | 0.106 | 50 |
|  | labs | EGFR | −0.36 | 9.53e-03 | 0.106 | 50 |
|  | labs | INTERLEUKIN6SERUM | −0.34 | 1.42e-02 | 0.123 | 50 |
|  | labs | RBC | 0.34 | 1.47e-02 | 0.123 | 50 |
|  | labs | MCV | 0.29 | 3.76e-02 | 0.206 | 50 |
|  | labs | VLDLCHOLESTEROLCAL | 0.29 | 3.84e-02 | 0.206 | 50 |
|  | nad_metabolites | 2PY | 0.29 | 3.99e-02 | 0.206 | 49 |

*Spearman rank correlations (ρ) between baseline cirNAD or icNAD and clinical endpoints. N = 50 (primary analysis cohort, both arms). Shown: endpoints with nominal p < 0.05. p (adj) = M_eff/Šidák corrected within the baseline stratum ( 135 tests).* *Bold: p (adj) < 0.05*
